# *Akkermansia muciniphila* secretome promotes α-synuclein aggregation in enteroendocrine cells

**DOI:** 10.1101/2021.02.12.430931

**Authors:** Dionísio Pedro Amorim Neto, Beatriz Pelegrini Bosque, João Vitor Pereira de Godoy, Paulla Vieira Rodrigues, Dario Donoso Meneses, Katiane Tostes, Celisa Caldana Costa Tonoli, Christian González-Billault, Matheus de Castro Fonseca

## Abstract

The notion that the gut microbiota play a role in neurodevelopment, behavior and outcome of neurodegenerative disorders is recently taking place. A number of studies have consistently reported a greater abundance of *Akkermansia muciniphila* in Parkinson’s disease (PD) fecal samples. Nevertheless, a functional link between *A.muciniphila* and sporadic PD remained unexplored. Here, we investigated whether *A.muciniphila* secretome could initiate the misfolding process of α-synuclein (αSyn) in enteroendocrine cells (EECs), which are part of the gut epithelium and possess many neuron-like properties. We found that *A*.*muciniphila* secretome is directly modulated by mucin, induces intracellular calcium (Ca^2+^) release, and causes increased mitochondrial Ca^2+^ uptake in EECs, which in turn leads to production of reactive oxygen species (ROS) and αSyn aggregation. However, these events were efficiently inhibited once we buffered mitochondrial Ca^2+^. Thereby, these molecular insights provided here offer evidence that bacterial secretome is capable of inducing αSyn aggregation in enteroendocrine cells.

**SYNOPSIS FIGURE DESCRIPTION:** The secretome isolated from the commensal gut bacterium *Akkermansia muciniphila* triggers intracellular Ca^2+^ signaling in enteroendocrine cells, leading to increased mitochondrial Ca^2+^ uptake. Mitochondrial Ca^2+^ overload leads to ROS generation culminating with αSyn phosphorylation and aggregation (left panel). All these events were inhibited once mitochondrial Ca^2+^ is buffered (right panel).

- Gram-negative gut bacterium *Akkermansia muciniphila* is consistently found more abundant in Parkinson’s disease patients.
- *Akkermansia muciniphila* protein secretome composition is directly modulated by mucin and induces an IP_3_-independent endoplasmic reticulum (ER)-calcium release in enteroendocrine cells.
- This Ca^2+^ release is triggered by direct activation of Ryanodine Receptors leading to increased mitochondrial Ca^2+^ uptake.
- Mitochondrial Ca^2+^ overload leads to ROS generation culminating with αSyn aggregation.
- Buffering mitochondrial Ca^2+^ efficiently inhibits *A.muciniphila-*induced αSyn aggregation in enteroendocrine cells.

## INTRODUCTION

Traditionally, Parkinson’s disease (PD) has been characterized as a progressive neurodegenerative disorder caused by loss of dopaminergic neurons in the *susbtantia nigra pars compacta* of the midbrain (Davie, 2008). Neuronal loss leads to Parkinsonism, an array of motor symptoms comprehending muscle rigidity, slowness, tremors and difficulty in controlling movement (Bernheimer et al, 1973). However, recent studies have shown that drug-naïve PD patients frequently report gastrointestinal complaints such as constipation and nausea and prolonged intestinal transit time even years before the disease is diagnosed (Adams-Carr et al, 2016; Martinez-Martin et al, 2011; Mun et al, 2016). Therefore, this pathology is now considered as a multisystemic disease gathering a plethora of non-motor symptoms (Chaudhuri et al, 2006; Greenland et al, 2019).

Currently, PD is the second most common neurodegenerative disease and the interest of the scientific community to unveil the cellular and molecular mechanisms of this complex pathology has grown substantially, triggered especially by the discovery of a number of causative monogenic mutations (Bekris et al, 2010). Nevertheless, these mutations only explain a small percentage of all PD cases since about 90% of cases are sporadic (de Lau & Breteler, 2006).

The key dogma of PD consists in the aggregation of the protein alpha-synuclein (αSyn) within neurons (Goedert et al, 2013; Spillantini et al, 1997). This presynaptic protein is linked genetically and neuropathologically to PD. It is accepted that αSyn aberrant soluble oligomeric conformations (protofibrils) are the toxic species that disrupt cellular homeostasis and lead to neuronal death through effects on several intracellular targets, including synaptic function (Stefanis, 2012). This aggregation process can be caused by genetic or sporadic factors due to mitochondrial dysfunction, oxidative stress and altered proteostasis (Greenamyre & Hastings, 2004). Although this toxic aggregation occurs more widely throughout the central system, abundant clinical and pathological evidence shows that misfolded αSyn is found in enteric nerves before it appears in the brain (Braak & Del Tredici, 2009; Braak et al, 2003a; Hawkes et al, 2010). It was recently reported that enteroendocrine cells (EECs), which are part of the gut epithelium and are directly exposed to the gut lumen and its microbiome, possess many neuron-like properties and connect to enteric nerves (Chandra et al, 2017). This leads to the hypothesis that PD might originate in the gut and then spread to the central nervous system via cell-to-cell prion-like propagation (Chandra et al, 2017). Such a concept has gathered significant momentum in recent years and great attention has been given to the brain-gut connection. Therefore, the gut microbiome raises as a promising target to be investigated in the outcome of sporadic PD. Several reports have shown that individuals with PD display an imbalanced gut microbiome (dysbiosis) (Heintz-Buschart et al, 2018; Hill-Burns et al, 2017; Keshavarzian et al, 2015) where commensal bacteria (e.g., phylum Firmicutes) are reduced, while pathogenic Gram-negative bacteria (*Proteobacteria* sp, *Enterobacteriaceae* sp, *Escherichia* sp.) and mucin-degrading Verrucomicrobiaceae, such as *Akkermansia muciniphila* are increased (Hill-Burns et al, 2017; Keshavarzian et al, 2015; Li et al, 2017; Scheperjans et al, 2015; Unger et al, 2016).

The mucin-degrading microorganism *A. muciniphila* (Derrien, 2004) comprehends about 1-4% of the fecal microbiome in humans (Naito et al. 2018). While numerous diseases have been associated with a decrease in *A. muciniphila* abundance (Grander et al, 2018; Schneeberger et al, 2015), an increase of this microorganism has been consistently reported in PD patients (Baldini et al, 2020). In addition, it was shown that *A. muciniphila* abundance had the largest contribution to the significantly altered metabolite secretion profiles of sporadic PD patients (Baldini et al, 2020).

The microbial surface and secreted proteins contain many proteins that interact with other microbes, host and/or environment (Tjalsma et al, 2000). Secretome proteins (e.g., receptors, transporters, adhesins, secreted enzymes, toxins) not only allow bacteria to interact with and adapt to their environment, but also modulate the host cells activities (Gagic et al, 2016). Identifying the effects of *A. muciniphila* protein secretome arising from the interaction of these secreted proteins with the enteroendocrine cells could therefore increase our current understanding on the cell mechanisms that could lead to one of the possible outcomes of sporadic PD.

Based on the common occurrence of gastrointestinal symptoms in PD, dysbiosis among PD patients, and strong evidence that the microbiota influences CNS function, in this work we investigated whether and how *A. muciniphila* conditioned media alters enteroendocrine cells homeostasis leading to αSyn aggregation. Herein, we found that *A. muciniphila* secretome induces an IP_3_-independent endoplasmic reticulum (ER)- calcium release by directly modulating Ryanodine Receptors (RYR), leading to increased mitochondrial calcium (Ca^2+^) uptake. Mitochondrial Ca^2+^ overload leads to ROS generation culminating with αSyn aggregation. In addition, these events were efficiently inhibited once we buffered mitochondrial Ca^2+^. These molecular insights provided here will push further the understanding of the pathogenesis of PD by offering mechanistic evidence that bacterial secretome is capable of inducing αSyn aggregation in enteroendocrine cells.

## RESULTS

### *Akkermansia muciniphila* growth curve pattern and secretome composition are modulated by mucin

*A. muciniphila* is a mucin-degrading Gram-negative bacterium of the phylum Verrucomicrobia (Derrien et al, 2004). However, the intestinal mucus layer is thought to be inversely correlated with *A. muciniphila* abundance in the gut (Sovran et al, 2019). Prolonged lack of dietary fibers induces damage to the mucus barrier and is directly associated with increased abundance of *A. muciniphila.* This would bring gut bacteria closer to the intestinal epithelium, which could trigger deleterious effects or other host compensatory responses (Desai et al, 2016). To test whether mucin could interfere with *A. muciniphila* secretome composition, the strain DSM-22959 was harvest and monitored for 72 hs in both BHI culture medium and BHI supplemented with 0.4% mucin (from porcine stomach, Type II) (Fig EV1). It is clearly observed that the addition of mucin maintains the growth of *A. muciniphila* in BHI medium (Fig EV1A). When mucin was provided, *A. muciniphila* grew faster at log phase and maintained a plateau for a longer time than when cultivated in mucin-free BHI medium. In addition, when analyzing the conditioned media (CM) by MS-MS mass spectrometry, we identified 350 differentially expressed proteins in the secretome obtained from *A. muciniphila* when cultivated for 36-40 hs (peak of growth for both conditions) in 0.4% mucin-supplemented BHI medium as opposed to only 86 in mucin-free medium (Fig EV1B). Therefore, the secretome of *A. muciniphila* is directly modulated by the presence of mucin.

### Intracellular calcium signaling is elicited by *Akkermansia muciniphila* mucin-free conditioned medium in a model of enteroendocrine cells

Enteroendocrine cells (EECs) are chemosensory cells distributed throughout all the mucosal lining of the intestine and with their apical surface exposed to the lumen of the organ. In addition, it was recently described that EECs also connect to enteric neurons (Bohorquez et al, 2015; Chandra et al, 2017; Liddle, 2018). Due to their location at the interface between gut contents and the nervous system, EECs provide a direct route for substances in the gut to affect neural function. The STC-1 cell line is widely accepted as a model of native EECs (McCarthy et al, 2015) due to the expression of several gastrointestinal hormones, including cholecystokinin (CCK) and peptide YY (PYY), whose secretion pattern is compared to that of native EECs (Hand et al, 2013; Hand et al, 2012; Sundaresan et al, 2013; Wang et al, 2002). Since native EECs are hard to culture or to be collected from intestinal tissue in a sufficient number for *in vitro* assays, STC-1 are considered an attractive cell model for evaluating properties of EECs.

Calcium (Ca^2+^) is known to regulate several important cell functions, such as secretion, proliferation, apoptosis, protein biosynthesis and folding (Alvarenga et al, 2016; Carafoli & Krebs, 2016; Fonseca et al, 2018; Guimaraes et al, 2017). In order to study the effects of *A. muciniphila* conditioned media in the fluctuations of intracellular Ca^2+^ signaling in STC-1 cells, we first stimulated Fluo-4/AM-loaded cells with 1 or 10% conditioned BHI medium (BHI CM) or unconditioned BHI medium (BHI). We observed that *A. muciniphila* mucin-free BHI CM induces a strong increase in Ca^2+^ transient in a concentration-dependent manner (Fig 1A-C). On the other hand, 0.4% mucin-supplemented BHI CM induced weaker Ca^2+^ signals when compared to the mucin-free condition (Fig EV2A-C). In order to observe whether this Ca^2+^ fluctuation was due to bacterial secretome and not to the unconditioned culture medium, STC-1 cells were also stimulated with mucin-supplemented or mucin-free unconditioned media (BHI) and no fluctuation on intracellular Ca^2+^ signals was observed (Fig EV3A-F).

**Fig 1.**
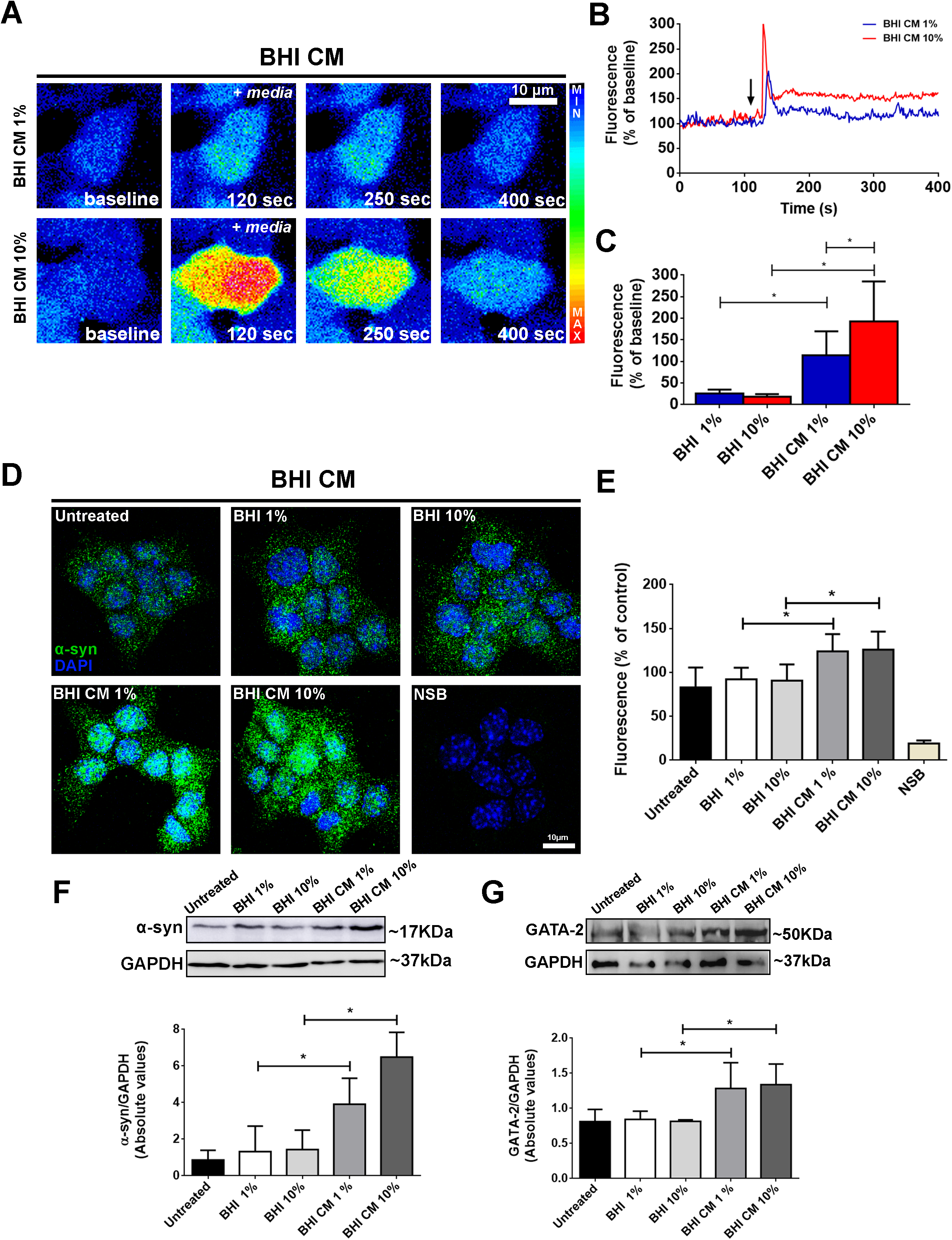
*Akkermansia muciniphila* conditioned medium induces intracellular calcium signals and increased levels of α-synuclein in STC-1 cells. **A.** Confocal microscopy imaging of STC-1 cells incubated with Fluo-4/AM (6μM) and stimulated with 1 or 10% *A. muciniphila* conditioned media (BHI CM) (scale bar: 10 μm). **B.** Representative time-course of total Ca^2+^ signal. Arrow indicates the moment when culture medium was applied. **C.** Quantification of the peak fluorescence following stimulation with 1 or 10% conditioned (BHI CM) and unconditioned (BHI) media. * P < 0.05 by two-way Student’s *t-test*. **D.** αSyn staining (green) in STC-1 cells after 48 hs incubation with 1-10% conditioned (BHI CM) or unconditioned media (BHI) demonstrating increased expression of the protein. Nuclei were stained with DAPI (blue) and immunofluorescence control is shown as NSB (non-specific binding control) (scale bar: 10 µm). **E.** Quantification of αSyn fluorescence intensity in images shown in (D). * P < 0.05 by two-way Student’s *t-test*. **F.** Immunoblots (upper image) of total cell lysates showing the increased expression of αSyn after 48 hours-incubation with 1-10% conditioned (BHI CM) or unconditioned media (BHI). Densitometric analysis shows increased expression of αSyn in 1-10% BHI CM condition when compared to 1-10% BHI. * P < 0.05 by two-way Student’s *t-test*. **G.** Immunoblots (upper image) of total cell lysates showing the increased expression of GATA-2 after 48 hours-incubation with 1-10% conditioned (BHI CM) or unconditioned media (BHI). Densitometric analysis shows increased expression of GATA-2 in 1-10% BHI CM condition when compared to 1-10% BHI. * P < 0.05 by two-way Student’s *t- test*. Data information: Data in (B) represent a representative tracing recorded from one individual STC-1 cell of each group. Data in (C and E-G) represent the mean ± SEM of three independent experiments. Whereas at least 55 individual cells were analyzed for calcium transient experiments and immunofluorescence, densitometric analysis of Western blot are derived from triplicates of three different experiments.

### Mucin-free *A. muciniphila* conditioned media increases expression of endogenous α-synuclein in STC- cells

Induced transient increase in free intracellular Ca^2+^ concentration by thapsigargin or Ca^2+^ ionophore chemical treatments lead to a significant increase in the number of cells presenting microscopically-visible αSyn aggregates (Nath et al, 2011). In addition, increased expression or decreased degradation of αSyn can initiate the formation of amyloid aggregates that can assemble to form Lewy bodies and Lewy neurites over the course of a lifetime (Hijaz & Volpicelli-Daley, 2020).

In addition, misfolded αSyn is found in enteric nerves before it appears in the brain (Braak & Del Tredici, 2009; Braak et al, 2003a; Hawkes et al, 2010). However, it is yet to be demonstrated whether the secretome of a gut bacterium could initiate this pathologic sequence of events.

Therefore, we next analyzed whether *A. muciniphila* CM could modulate αSyn homeostasis in STC-1 cells. MTS assay confirmed that 48h-incubation of cells with 1 or 10% BHI CM or BHI did not decrease cell viability. In addition, secretion of CCK was not altered (Fig EV4A and B). When STC-1 cells were incubated with 1 or 10% mucin-free BHI CM for 48 hs, but not with the unconditioned one (BHI), we detected a significant clear overexpression of αSyn analyzed by immunofluorescence and Western blotting (Fig 1D-F). However, this was not observed when the cells were incubated with 0.4% mucin-containing BHI CM (Fig EV2D-F).

The SNCA gene expression in neurons, which encodes for αSyn, is known to be controlled by the GATA-2 transcription factor (Scherzer et al, 2008), which also plays a crucial role in central nervous system development, and erythroid cells differentiation (Nardelli et al, 1999). In addition, GATA-2 has a critical role in neuronal development, particularly in cell fate specification of catecholaminergic sympathetic neurons (Bilodeau et al, 2001; Tsarovina et al, 2004). We observed that STC-1 cells not only express GATA-2 transcription factor but also exhibit increased expression of this factor when incubated with either 1% or 10% mucin-free *A. muciniphila* BHI CM. This supports the idea that *A .muciniphila* secretome upregulates GATA-2 which in turn induces SNCA overexpression (Fig 1G).

In order to confirm if these observed effects were specifically due to *A. muciniphila* conditioned medium, we conducted the same set of above experiments employing *Escherichia coli* (*E. coli*) conditioned medium. *E. coli* was chosen because it is an abundant Gram-negative microorganism from the gut. This strain was also cultivated in BHI medium under anaerobic condition as for *A. muciniphila*. Although we also observed a transient increase in free intracellular Ca^2+^ in STC-1 cells stimulated with 1 or 10% *E. coli* BHI CM, the amplitude of the signal was smaller than the one elicited by *A. muciniphila,* as shown in Fig EV5A-E. In addition, we did not detect alteration on αSyn expression levels when STC-1 cells were incubated for 48 hs with *E. coli* CM (Fig EV5F-H).

In summary, *A. muciniphila* mucin-free CM leads to a transient increase in free intracellular Ca^2+^ and induces GATA2-regulated-overexpression of αSyn in the STC-1 enteroendocrine cell model.

### *A. muciniphila* conditioned medium induces calcium release from stores in the endoplasmic reticulum in an IP_3_-independent manner

Several maneuvers were performed to define the mechanism by which *A. muciniphila* mucin-free CM increases free cytoplasmic Ca^2+^ in STC-1 cells. To determine the source of the Ca^2+^, cells were stimulated in Ca^2+^-free medium. We observed that *A .muciniphila* CM induced cytoplasmic Ca^2+^ oscillations in a concentration-dependent manner even in Ca^2+^-free medium (Fig 2A and B). Additionally, induced-Ca^2+^ signals initiate/predominate in the cytoplasm (Fig EV6) and were elicited in a similar fraction of STC-1 cells regardless of the presence of extracellular Ca^2+^. On the other hand, selective depletion of stored calcium by 10µM thapsigargin significantly blocked Ca^2+^ oscillations-induced by *A. muciniphila* CM (Fig 2C and D). Thereby, these findings demonstrate that *A. muciniphila* CM sample increases cytoplasmic Ca^2+^ levels by mobilizing intracellular Ca^2+^ stores.

**Fig 2.**
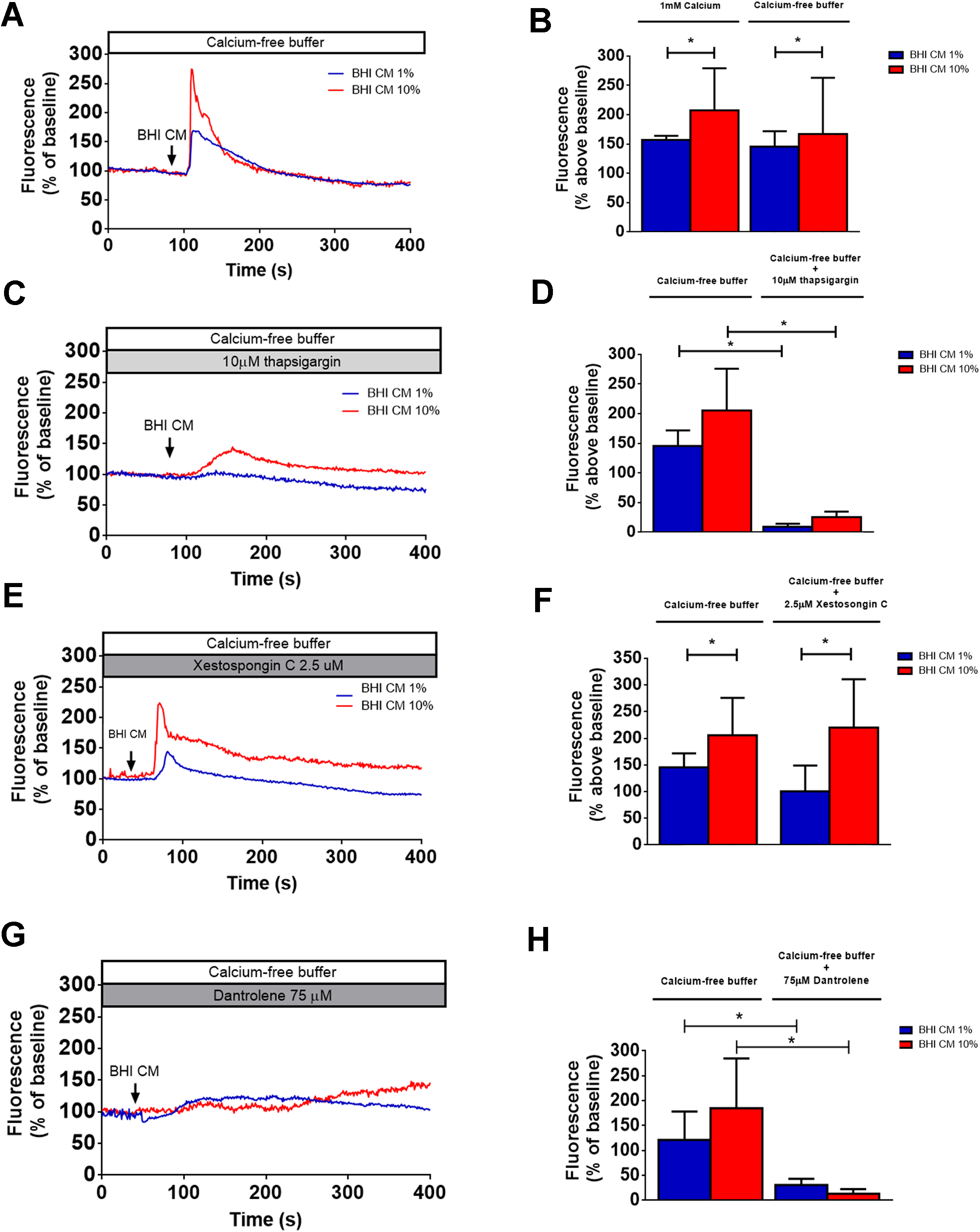
*Akkermansia muciniphila* conditioned medium induces InsP3-independent intracellular calcium signals by acting directly on ryanodine. **A.** STC-1 cells were stimulated with 1 or 10% *A. muciniphila* conditioned media (BHI CM) in the presence of Ca^2+^-free buffer. Graph shows a representative time-course of total Ca^2+^ signal in STC-1cells. The arrow indicates the time when culture medium was applied. **B.** Quantification of the peak fluorescence following cells stimulation with 1 or 10% conditioned (BHI CM) and unconditioned (BHI) media in the presence of 1 mM Ca^2+^ - buffer or Ca^2+^ - free buffer. * P < 0.05 by two-way Student’s *t-test*. **C.** STC-1 cells were incubated with 10 µM thapsigargin for 30 min and stimulated with 1 or 10% *A. muciniphila* conditioned media (BHI CM) in the presence of Ca^2+^-free buffer containing 10 µM thapsigargin. Graph shows a representative time-course of total Ca^2+^ signal in STC-1cells. Arrow indicates the time when culture medium was applied. **D.** Quantification of the peak fluorescence following cells stimulation with 1 or 10% conditioned (BHI CM) and unconditioned (BHI) media shows that the Ca^2+^ signal induced by BHI CM is blocked by thapsigargin 10 µM. * P < 0.001 by two-way Student’s *t-test*. **E.** STC-1 cells were incubated with 2.5 µM xestospongin C for 30 min and stimulated with 1 or 10% *A. muciniphila* conditioned media (BHI CM) in the presence of Ca^2+^-free buffer containing 2.5 µM xestospongin C. Graph shows a representative time-course of total Ca^2+^ signal in STC-1 cells. The arrow indicates the time when culture medium was applied. **F.** Quantification of the peak fluorescence following cells stimulation with 1 or 10% conditioned (BHI CM) and unconditioned (BHI) media shows that the Ca^2+^ signal induced by BHI CM is not blocked by the InsP3 receptor inhibitor xestospongin C (2.5 μM). * P < 0.05 by two-way Student’s *t-test*. **G.** STC-1 cells were incubated with 75 µM dantrolene for 30 min and stimulated with 1 or 10% *A. muciniphila* conditioned media (BHI CM) in the presence of Ca^2+^-free buffer containing 75µM dantrolene. Graph shows a representative time-course of total Ca^2+^ signal in STC-1cells. The arrow indicates the time when culture medium was applied. **H.** Quantification of the peak fluorescence following cells stimulation with 1 or 10% conditioned (BHI CM) and unconditioned (BHI) media shows that the Ca^2+^ signal induced by BHI CM is completely blocked by the RYR receptor inhibitor, dantrolene (75 μM). (n = 3 individual experiments, *p<0.05, Bonferroni post-tests. Data are expressed as media ± SEM). * P < 0.001 by two-way Student’s *t-test*. Data information: Data in (A, C, E and G) represent a representative tracing recorded from one individual STC-1 cell of each group. Data in (B, D, F and H) represent the mean ± SEM of three independent experiments in which at least 55 individual cells were analyzed for calcium transient.

A classic manner by which extracellular factors initiate an intracellular Ca^2+^ mobilization is by generating InsP3 to bind and release Ca^2+^ from InsP3 receptors in the endoplasmic reticulum (Divecha et al, 1991). In order to investigate whether the cytoplasmic Ca^2+^ increase was triggered by InsP3 generation, we stimulated STC-1 cells in the presence of the InsP3 receptor inhibitor xestospongin C (Gafni et al, 1997). Incubation of cells for 30 min and continuous perfusion with 2.5 µM xestospongin C did not impair *A. muciniphila* CM -induced Ca^2+^ mobilization, suggesting an InsP3- independent release of intracellular Ca^2+^ stores (Fig 2E and F). To go further into the mechanism by how *A. muciniphila* CM evokes Ca^2+^ release from intracellular stores, we incubated cells for 30 min with dantrolene (75 μM), an inhibitor of Ca^2+^ release through ryanodine receptor (RYR) channels (Hainaut & Desmedt, 1974; Morgan & Bryant, 1977; Zhao et al, 2001). In the presence of 75 μM dantrolene, only a very small Ca^2+^ increase was observed following stimulation with 1 or 10% *A. muciniphila* CM (Fig 2G and H).

Thus, dantrolene eliminated *A. muciniphila* CM Ca^2+^ response in enteroendocrine cells. Taken together, these results show that *A. muciniphila* secretome works as a physiological RYR gating agent, eliciting intracellular Ca^2+^ signals by directly modulating RYR in the cytoplasm.

### Mitochondrial calcium overload, senescence and intracellular ROS production is elicited by *A. muciniphila* secretome

Global changes in Ca^2+^ homeostasis accompanied by the alteration in cellular bioenergetics status and thereby imposing oxidative stress in cells are reported in PD (Ludtmann & Abramov, 2018; Surmeier & Schumacker, 2013). The cytosolic Ca^2+^ concentration in unstimulated cells is maintained at low levels (∼100 nM) by several enzymes that translocate Ca^2+^ ions into intracellular stores or across the plasma membrane. Moreover, Ca^2+^ uptake into the mitochondria is not limited to the control of organelle function, but also has a direct impact on the intracellular Ca^2+^ signals evoked by agonist stimulation in the cytosol through modulation of their kinetics and spatial dimensions (Tinel et al, 1999). Enhanced cytosolic Ca^2+^ concentration, on the other hand, affects the bioenergetics of the cells by promoting increased ATP demand (Hill et al, 2012). Furthermore, this alteration in cytosolic Ca^2+^ hampers the normal Ca^2+^ handling by various intracellular organelles, including mitochondria, and threatens neuronal survival. Although well established for neuronal cells, there is still a gap regarding changes in mitochondrial Ca^2+^ dynamics in enteroendocrine cells due to gut microbiome stimulation and how this event might be related to αSyn homeostasis.

Thereby, we aimed at evaluating mitochondrial Ca^2+^ under stimulation with *A. muciniphila* secretome. When STC-1 cells loaded with the mitochondrial Ca^2+^ indicator Rhod-2/AM dye were stimulated with 10% of *A. muciniphila* CM, we observed a significant increase in mitochondrial Ca^2+^ uptake when compared to unconditioned BHI medium (Fig 3A and B). In addition, when we incubated the cells for 48 hs with 1 or 10% CM and stimulated with ATP (10 µM), mitochondrial fluorescence was dramatically increased in the group incubated with 10% BHI CM when compared to 1% BHI CM or unconditioned BHI medium (1 and 10%) suggesting that long exposure to *A. muciniphila* secretome induces increased uptake of Ca^2+^ by the mitochondria (Fig 3C and D).

**Fig 3.**
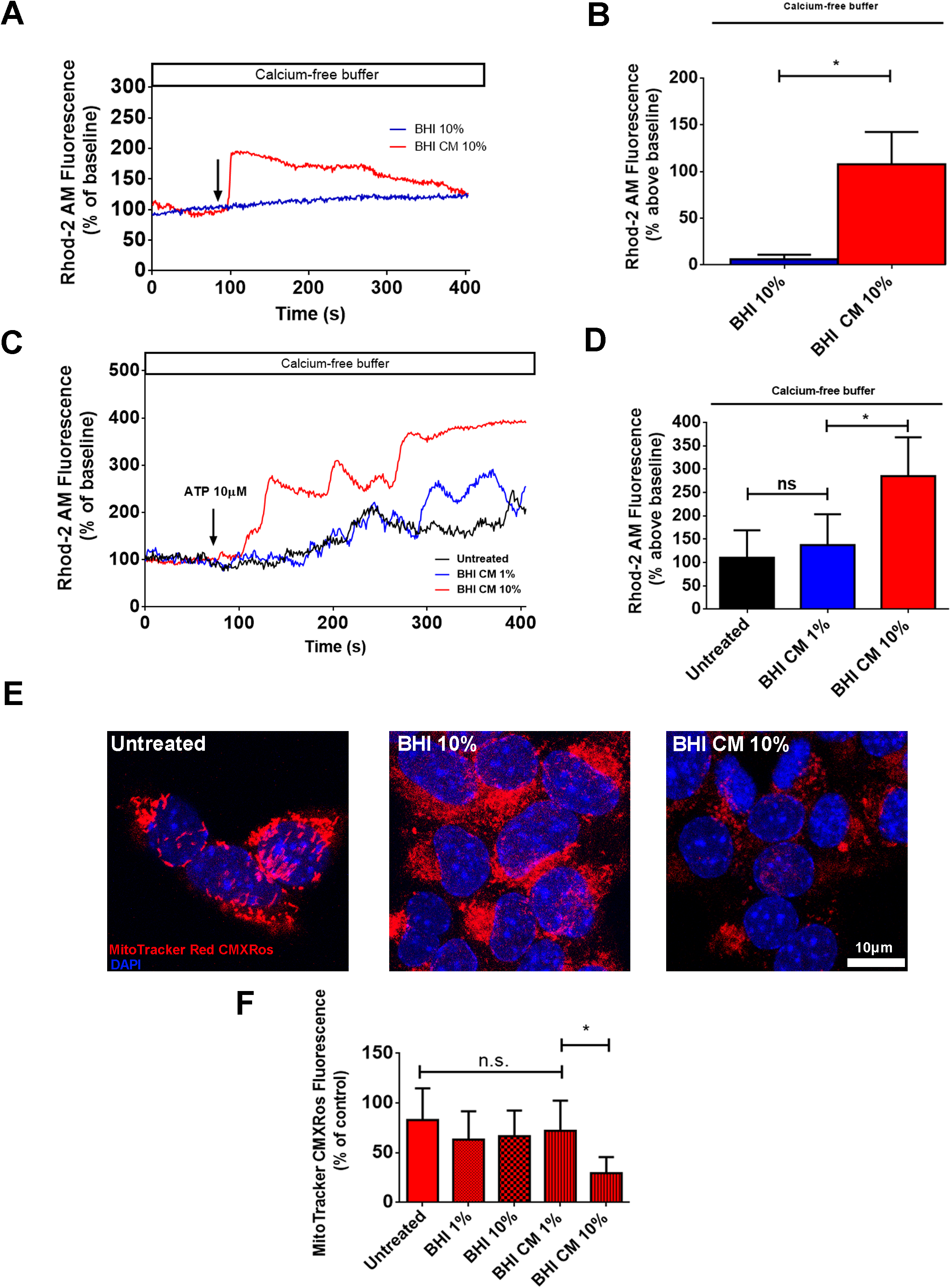
Increased mitochondrial Ca^2+^ uptake elicited by *Akkermansia muciniphila* conditioned media leads to mitochondrial stress and reduced **Δ**Ψm. **A.** Representative time-course of mitochondrial Ca^2+^ signal. Cells were incubated with the mitochondrial Ca^2+^ indicator Rhod-2/AM and stimulated with 10% *A. muciniphila* conditioned medium (BHI CM) in the presence of Ca^2+^-free buffer. The arrow indicates the time when culture medium was applied. **B.** Graphs show quantification of the peak of fluorescence following stimulation with 10% BHI CM. * P < 0.001 by two-way Student’s *t-test*. **C.** Representative time-course of mitochondrial Ca^2+^ signal of cells incubated for 48 hs with 1-10% *A. muciniphila* conditioned medium (BHI CM) and stimulated with 10 µM ATP in the presence of Ca^2+^-free buffer. The arrow indicates the moment when culture medium was applied. **D.** Graphs show quantification of the peak of fluorescence following stimulation with ATP. * P < 0.05; Ns, not significant by two-way Student’s *t-test*. **E.** Confocal images of STC-1 cells incubated for 48 hs with 10% BHI CM and then stained with MitoTracker Red CMXRos (red). Nuclei were stained with DAPI (blue). **F.** Quantification of fluorescent signal in untreated and treated cells. * P < 0.05; Ns, not significant by two-way Student’s *t-test*. Data information: Data in (A and C) represent a representative tracing recorded from one individual STC-1 cell of each group. Data in (B, D and F) represent the mean ± SEM of three independent experiments in which at least 55 individual cells were analyzed.

As previously mentioned, enhanced, or sustained Ca^2+^ stress results in mitochondrial injury due to Ca^2+^ overload. Excessive mitochondrial Ca^2+^ uptake or impaired Ca^2+^ efflux influences mitochondrial membrane potential (ΔΨm) leading to depolarization of mitochondrial inner membrane, swelling of the organelle, and ultimately cell death (Calvo-Rodriguez et al, 2020; Di Lisa & Bernardi, 2009; Gunter et al, 1998; Williams et al, 2013). In order to observe whether mitochondrial Ca^2+^ uptake induced by *A. muciniphila* CM could lead to mitochondrial damage, we monitored ΔΨm in STC-1 cells under *A. muciniphila* CM incubation for 48 hs. After treatment, cells were stained with the mitochondrial-targeted probe Mitotracker Red CMXRos, which accumulates in mitochondria depending on its membrane potential and has been widely used as an indicator of reduced ΔΨm (Jacotot et al, 2000a; Jacotot et al, 2000b). As can be observed on Fig 3E and F, cells incubated with 10% CM presented a reduced fluorescent signal of the probe what suggests impaired membrane potential. In addition, mitochondria displayed more fragmented and rounded, showing a swelled morphology, which was not observed when cells were incubated either with 1% CM or with unconditioned BHI medium.

Altogether, the results described so far demonstrate that *A. muciniphila* secretome induces exacerbated mitochondrial Ca^2+^ uptake, which in turn is the driven force that causes mitochondrial damage, reflected by a loss of membrane ΔΨm.

### Increased intracellular ROS level, α-synuclein phosphorylation and aggregation as a consequent event of *A. muciniphila* secretome stimulation of enteroendocrine cells

It is suggested that endogenous ROS mainly modulate cell signaling locally and stimuli that promote ROS formation or mitochondrial alterations highly correlate with mutant αSyn phosphorylation at Serine 129 (Ser129), a promoter of αSyn aggregation propensity and toxicity in PD (Karampetsou et al, 2017; Perfeito et al, 2014; Tenreiro et al, 2014). Therefore, we next measured intracellular levels of ROS under stimulation with 1 or 10% *A. muciniphila* CM by live cell imaging. STC-1 cells were incubated for 30 min with DHE and continuously perfused with buffer containing 1 or 10% CM. Buffer/unconditioned media and H_2_O_2_ (100 µM) perfusion were used as negative and positive controls, respectively. The real-time fluorescence measurement indicates that the surge of ROS level after H_2_O_2_ or 1-10% CM stimulation was significantly higher than stimulation with buffer or 1-10% unconditioned BHI media for 5 min (Fig 4A and B). In addition, cells stimulated with 1 or 10% CM presented increased DHE fluorescence in a similar manner.

**Fig 4.**
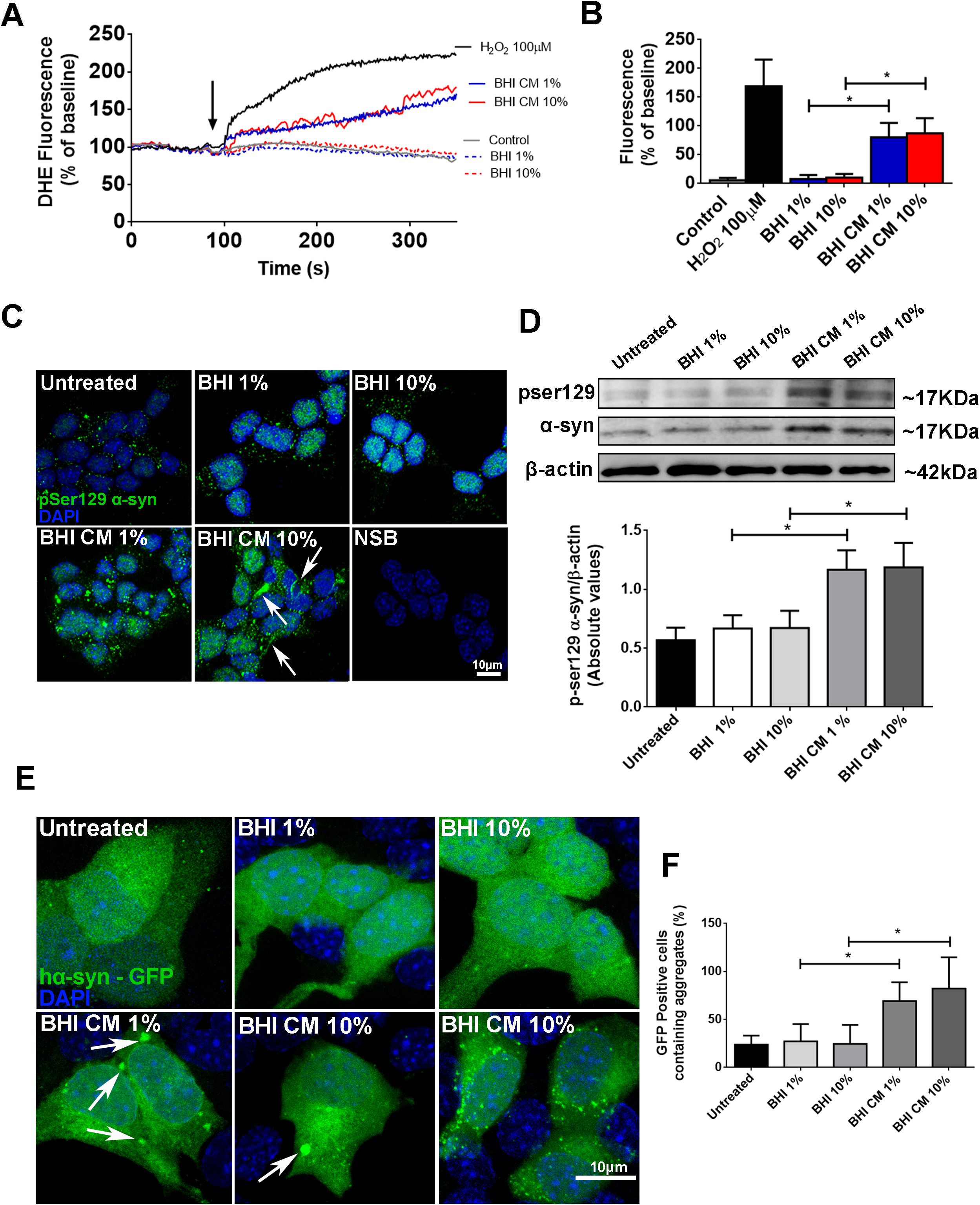
α-synuclein phosphorylation and aggregation as a result of increased intracellular levels of ROS due to *Akkermansia muciniphila* conditioned media treatment of enteroendocrine cells. **A.** Time lapse of ROS production in STC-1 cells measured by DHE fluorescence intensity under confocal live imaging. 1 and 10 % BHI or BHI CM was used as stimuli. 100 mM H_2_O_2_ was used as positive control. **B.** Quantitative summary of the effects of *A. muciniphila* conditioned and unconditioned media on ROS production. * P < 0.001 by two-way Student’s *t-test*. **C.** Confocal images of pSer129 αSyn staining (green) in STC-1 cells after 48 hs incubation with 1-10% conditioned (BHI CM) or unconditioned media (BHI) demonstrating increased phosphorylation of the protein. Nuclei were stained with DAPI (blue) and immunofluorescence control is shown as NSB (non-specific binding control). Arrows point to aberrant fibrillary-like structures. (Scale bar: 10 µm). **D.** Immunoblots (upper image) of total cell lysates showing the increased phosphorylation of αSyn on Ser129 (normalized against total αSyn) after 48 hours-incubation with 1/10% conditioned (BHI CM) or unconditioned media (BHI). Densitometric analysis shows phosphorylation of αSyn (pSer129) in 1-10% BHI CM condition when compared to 1/10% BHI. * P < 0.05 by two-way Student’s *t-test*. **E.** Confocal images of hαSyn GFP-tagged plasmid transfected into STCI-1 cells exhibits diffuse distribution in untreated or BHI-treated cells. Cells treated with 1-10% BHI CM medium forms inclusions of different sizes (bottom images). (Scale bar: 10 µm). **F.** Graph shows the number of GFP-positive cells containing inclusions in each condition. * P < 0.05 by two-way Student’s *t-test*. Data information: Data in (A) represent a representative tracing recorded from one individual STC-1 cell of each group. Data in (B and F) represent the mean ± SEM of three independent experiments in which at least 55 individual cells were analyzed. Densitometric analysis of western blot (D) are derived from triplicates of three different experiments.

As mentioned, stimuli that promote intracellular ROS formation and mitochondrial damage highly correlate with αSyn phosphorylation at Ser129, an event that may precede cell degeneration in PD (Perfeito et al, 2014). Previous observations have shown that both nigral and dorsal motor nucleus of the vagus nerve neurons present a high vulnerability to oxidative challenges (Musgrove et al, 2019). Since the nigro-vagal pathway that controls gastric tone and motility connect these brain regions, it raises the possibility that an oxidative injury may be relayed and possibly amplified through this anatomical and functional connection.

In order to evaluate whether increases ROS levels induced by *A. muciniphila* CM could promote αSyn phosphorylation and aggregation, we incubated the cells for 48 hs in the presence of CM or unconditioned BHI media and directed them to immunofluorescence and Western blotting. Confocal microscopy images showed strong deposits of pSer129-αSyn in STC-1 cells incubated with 1 and 10% CM (Fig 4C). In addition, quantification by Western blotting showed a 2-3 fold increase of p-Ser129- αSyn in cells treated with the secretome when normalized against total αSyn (Fig 4D). To establish whether *A. muciniphila* CM-induced p-Ser129 αSyn might play a role on αSyn aggregation in our STC-1 cell model, we transfected cells with full-length human αSyn-GFP-tagged and incubated them with unconditioned or CM for 48hs. Unconditioned BHI media (1 or 10%) did not cause αSyn to form cellular inclusions. However, 1 and 10% CM led to the formation of small to large αSyn granules within the cytoplasm (Fig 4E). When we quantified the number of GFP-positive cells containing intracellular aggregates, we observed that over 50% of the cells stimulated with *A. muciniphila* CM contained αSyn granules (Fig 4F). Thereby, *A. muciniphila* secretome induces intracellular αSyn aggregation in enteroendocrine cell model.

### Mitochondrial calcium buffering reverts the damaging effects to mitochondria and prevents α-synuclein aggregation

Inhibition of mitochondrial Ca^2+^ uptake was shown to diminish the oxidative stress in substantia nigra *pars compacta* dopaminergic neurons (SNpc DNs) suggesting that mitochondrial oxidative stress could also be due to mitochondrial Ca^2+^ overload (Guzman et al, 2010). Several lines of investigation point out to mitochondrial Ca^2+^ imbalance as key factor to be modulated in order to control the progression of PD. In order to observe whether modulating mitochondrial Ca^2+^ in enteroendocrine cells could reverse intracellular ROS generation and αSyn aggregation, we transfected the cells with parvalbumin (PV) fused to a mitochondrial targeting sequence (MTS) and GFP (Guerra et al, 2011). Parvalbumin (PV) is a cytosolic Ca^2+^-binding protein of the large EF-hand protein family, involved in intracellular Ca^2+^ regulation and buffering. GFP targeted to the mitochondrial matrix was used as a control (MTS-GFP) (Fig 5A).

**Fig 5.**
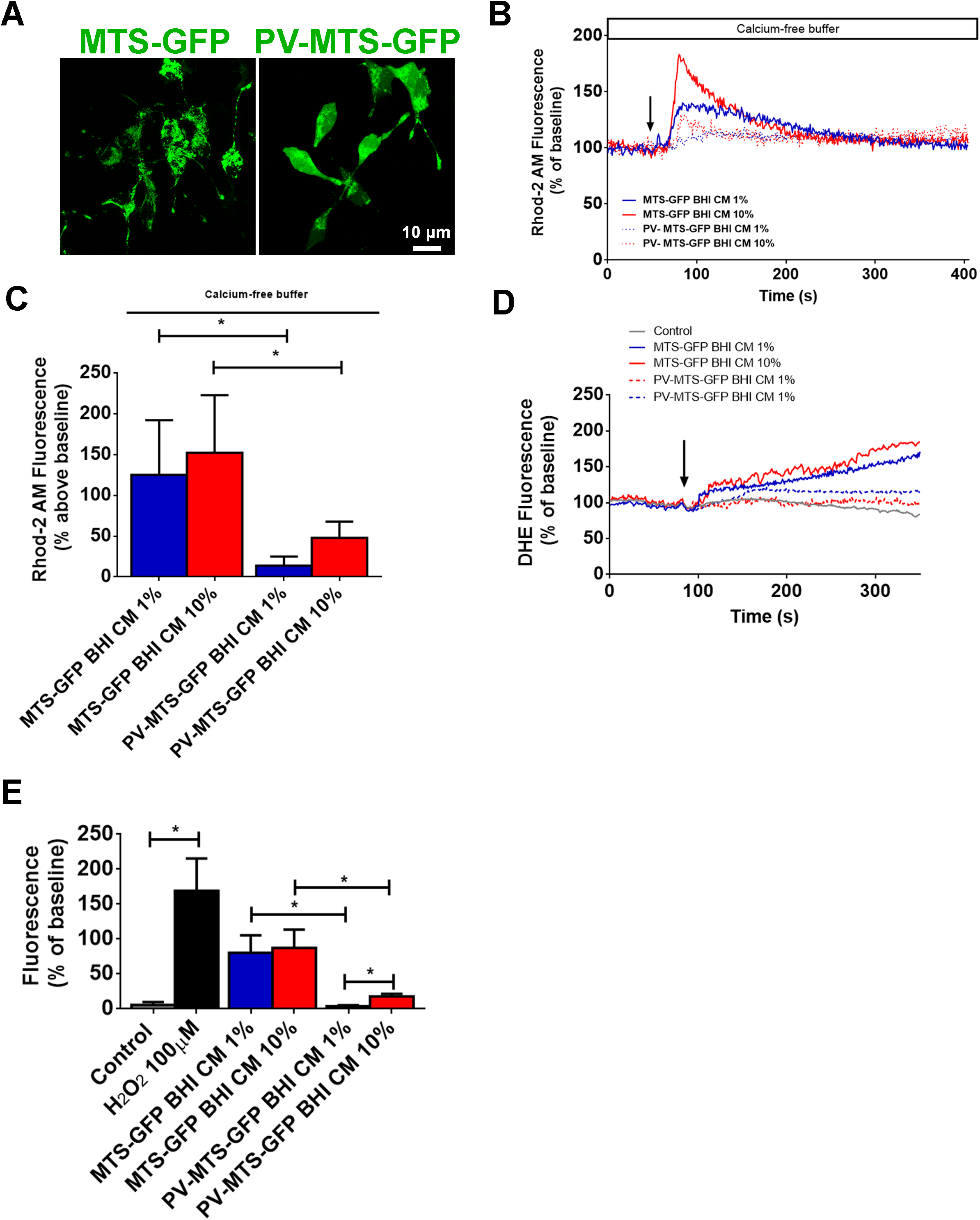
Mitochondrial Ca^2+^ buffering reduces intracellular ROS levels elicited by *Akkermansia muciniphila* conditioned medium. **A.** Confocal images of STC-1 cells transfected with mitochondrial parvalbumin (PV) expression and control vectors showing the expression and mitochondrial localization of targeted PV-MTS-GFP and MTS-GFP fusion proteins. Scale bar= 10 µM. **B.** Representative changes in mitochondrial Ca^2+^ signals over time are shown. Cells transfected with the indicated vectors were loaded with Rhod-2/AM and induced by 1- 10% unconditioned (BHI) or conditioned medium (BHI-CM) (arrow). Ca^2+^ signals were attenuated in cells expressing PV in mitochondria. **C.** Peak Ca^2+^ signals were observed in three separate experiments for STC- cells transfected with MTS-GFP, and cells transfected with PV-MTS-GFP. * P < 0.05 by two-way Student’s *t-test*. **D.** Representative changes in intracellular ROS levels over time are shown. Cells transfected with the indicated vectors were loaded with DHE and induced by 1/10% unconditioned (BHI) or conditioned medium (BHI-CM) (arrow). DHE fluorescence intensity was significantly reduced in cells expressing PV-MTS-GFP fusion protein. **E.** Peak ROS signals were observed in three separate experiments for STC- cells transfected with MTS-GFP, and cells transfected with PV-MTS-GFP stimulated with each represented condition. * P < 0.05 by two-way Student’s *t-test*. Data information: Data in (B and D) represent a representative tracing recorded from one individual STC-1 cell of each group. Data in (C and E) represent the mean ± SEM of three independent experiments in which at least 55 individual cells were analyzed.

One or 10% BHI CM elicited a robust increase in mitochondrial Ca^2+^ in cells expressing MTS-GFP alone, but this was reduced by approximately 90% in cells expressing PV in mitochondria (in Fig 5B and C). These results demonstrated that PV- MTS-GFP was correctly targeted to the mitochondrial matrix and efficiently buffered mitochondrial Ca^2+^ overload driven by stimulation with *A. muciniphila* conditioned medium.

Once mitochondrial Ca^2+^ was buffered, the next set of experiments aimed to observe whether the damaging effects caused by *A. muciniphila* conditioned medium could be prevented. When we stimulated the cells expressing PV-MTS construct with 1 and 10% BHI CM, the increase in intracellular ROS was significantly suppressed (Fig 5D and E) indicating that mitochondrial Ca^2+^ buffering prevents intracellular oxidative stress.

To test the effect of mitochondrial Ca^2+^ on Ser129-phosphorylation of αSyn induced by *A. muciniphila* conditioned medium, we incubated the transfected cells with 1 or 10% CM for 48hrs. Total cell lysate evaluated by Western blotting showed that levels of Ser129-phosphorylated αSyn significantly decreased in PV-MTS expressing cells when compared with control cells (MTS-GFP) (Fig 6A and B). However, no effect was observed in the total expression level of αSyn, which remained higher when compared to untreated cells (Fig 6A and C).

**Fig 6.**
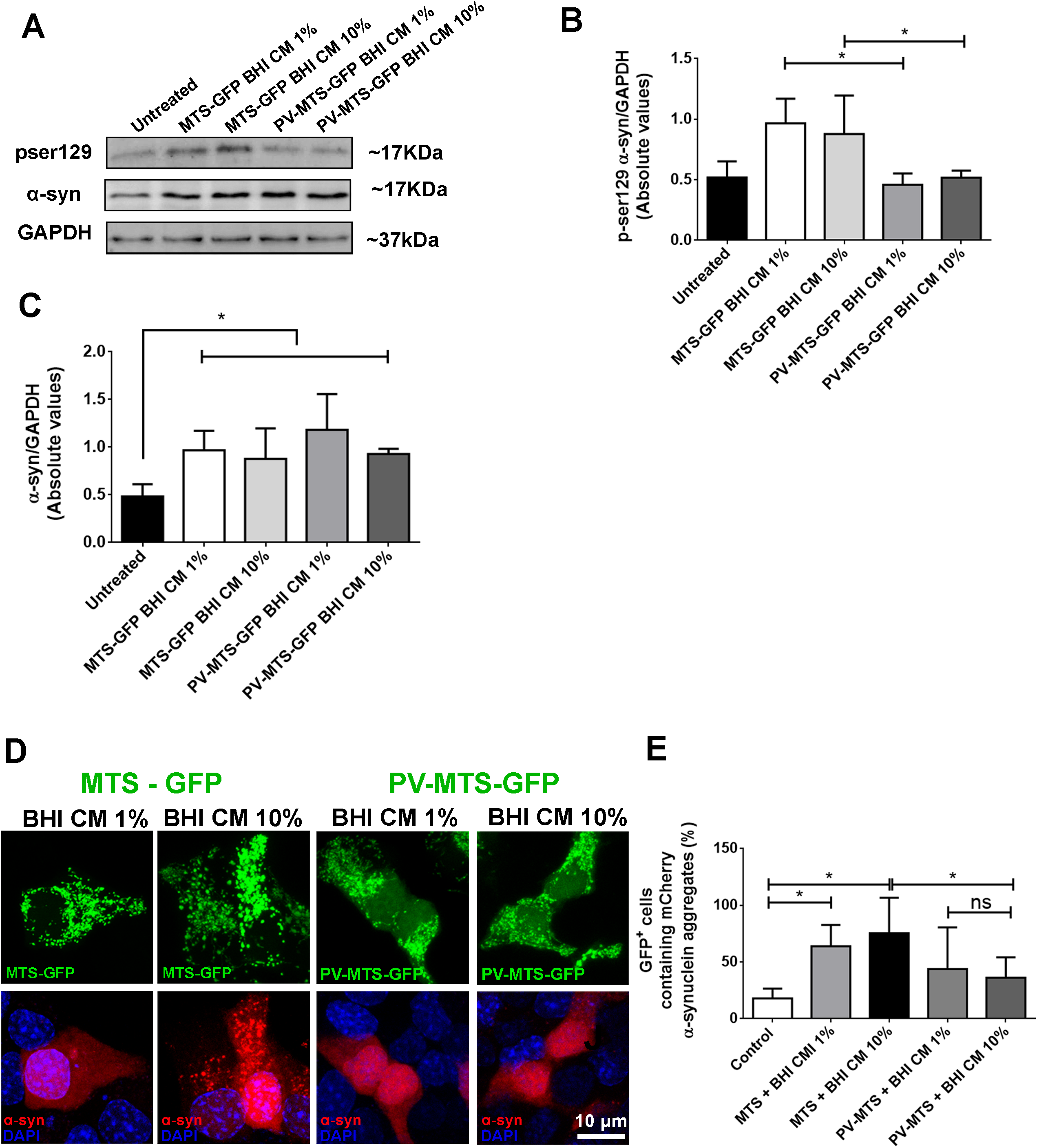
α-synuclein phosphorylation and aggregation induced by *Akkermansia muciniphila* conditioned medium are prevented due to mitochondrial Ca^2+^ buffering. **A.** Immunoblots (upper image) of total cell lysates from PV-MTS-GFP or MTS-GFP transfected cells showing a decrease in αSyn phosphorylation on Ser129 after 48 hours-incubation with 1-10% conditioned (BHI CM) or unconditioned media (BHI). **B.** Densitometric analysis shows that mitochondrial Ca^2+^ buffering reduced αSyn phosphorylation in 1-10% BHI CM condition when compared to cells expressing MTS-GFP fusion protein. * P < 0.05 by two-way Student’s *t-test*. **C.** Densitometric analysis shows that mitochondrial Ca^2+^ buffering did not reduce αSyn expression induced by 1/10% BHI CM condition when compared to cells expressing MTS-GFP fusion protein. * P < 0.05 by two-way Student’s *t-test*. **D.** Confocal images of STC-1 cells co-transfected with PV-MTS-GFP and αSyn- mCherry tagged construct show reduced number of intracellular αSyn aggregates after 48 hs incubation with 1-10% BHI CM. (scale bar: 10µM) **E.** Quantification of the number of GFP-positive cells containing mCherry-tagged αSyn aggregates. Data are expressed as percentage of total GFP-positive cells per image. * P < 0.05; Ns, not significant by two-way Student’s *t-test*. Data information: Densitometric analysis of western blot (B and C) are derived from triplicates of three different experiments. Data in (E) represent the mean ± SEM of three independent experiments in which at least 55 individual cells were analyzed.

We then extended our observation that mitochondrial Ca^2+^ can suppress intracellular ROS generation and αSyn phosphorylation to the formation of αSyn aggregates. Hence, we double-transfected cells with the PV-MTS-GFP construct and human αSyn mCherry-tagged. Large number of αSyn aggregates were observed in cells expressing the control construct (MTS-GFP) after 48 hs of treatment with 1 or 10% conditioned medium. However, the number of αSyn aggregates in cells expressing the PV-MTS-GFP constructed was markedly reduced (Fig 6D and E).

Taking together, these findings provide evidence on the mechanism by which *A. muciniphila* conditioned media induces αSyn aggregation in this enteroendocrine cell line.

## DISCUSSION

Parkinson’s disease is a growing health concern for an ever-aging population. Although genetic risks have been identified, environmental influences and gene-environment interactions are so far considered responsible for most PD cases (Nalls et al., 2014, Ritz et al., 2016). Besides the plethora of neurological and motor symptoms, PD patients present prominent gut manifestations (Abbott et al, 2001; Adams-Carr et al, 2016; Cersosimo et al, 2013; Chaudhuri et al, 2006; Jost, 2010; Klingelhoefer & Reichmann, 2015; Mertsalmi et al, 2017).

Aggregates of the protein aSyn is a hallmark of PD. Interestingly, aSyn pathology in PD is not limited to the brain. It was also observed in the peripheral nervous system, including the enteric nervous system (Wakabayashi et al, 2010). Therefore, the interaction between the gut microbiota, EECs and αSyn aggregation in PD is receiving increasing attention. The idea that aSyn aggregation process is initiated in the gut following continuous gastrointestinal symptoms aggravation and spreading to the nervous system in a prion-like manner has gathered significant force in recent years. Some pathophysiological evidence helps to support this notion: αSyn inclusions appear earlier in the enteric nervous system and the glossopharyngeal and vagal nerves (Braak et al., 2003, Shannon et al., 2012); and vagotomized individuals are at reduced risk for PD (Svensson et al., 2015). In addition, injection of αSyn fibrils into the gut tissue of healthy rodents seemed sufficient to induce disease within the vagus nerve and brainstem (Kim et al, 2019).

The finding that EECs connect to nerves raises an array of possibilities for how nutrients, bacteria, toxins, and potential pathogens gain access to and communicate with the nervous system. The discovery of αSyn in EECs, which are directly exposed to *A. muciniphila* secretome in the gut lumen and connected to enteric nerves, provides a location in which misassemble and spread of α-synuclein could initiate (Chandra et al, 2017). However, the knowledge of how αSyn aggregation initiates in the gut and spreads to the central nervous system via retrograde transmission and whether the gut microbiome could directly trigger this process remains controversial (Burke et al., 2008).

The specific mechanisms by which gut bacteria promote αSyn-mediated pathophysiology are likely diverse, complex, and poorly explored. Nonetheless, in this work we have identified that mitochondrial Ca^2+^ overload in EECs led by the secretome of a commensal gut bacterium *A. muciniphila* is a molecular pathway by which αSyn homeostasis is disturbed in EECs, providing experimental support for a gut-microbial connection to PD.

Since its discovery in 2004, *Akkermansia muciniphila* (Derrien et al, 2004) has garnered a great amount of scientific attention. It has been shown that intestinal *Akkermansia* abundance is significantly reduced in a many metabolic disorders, including type 2 diabetes, obesity, dyslipidemia (Derrien et al, 2017). Therefore, this has stimulated several studies in order to investigate *Akkermansia* supplementation. Evidence shows *Akkermansia* supplementation restores epithelial mucosal integrity, reduces weight gain and fat accumulation in the liver, improves glucose tolerance, and reduces inflammation and metabolic endotoxemia in animal models of diabetes and obesity (Derrien et al., 2017; Plovier et al., 2017).

However, we cannot rule out recent reports showing that alteration in gut microbiota is associated with PD. Several lines of evidence suggest that PD patients present strong gut dysbiosis with remarkable abundance of *A. muciniphila* which is consistently high in PD stool samples (Baldini et al, 2020; Bedarf et al, 2017; Desai et al, 2016; Heintz-Buschart et al, 2018; Hill-Burns et al, 2017; Li et al, 2017; Lin et al, 2019; Mertsalmi et al, 2017; Nishiwaki et al, 2020; Unger et al, 2016; Vidal-Martinez et al, 2020). Additionally, emerging studies have described increased abundance of *A. muciniphila* in multiple sclerosis (MS), one of which enrolled monozygotic twins discordant for MS diagnosis, therefore excluding genetic confounders (Berer et al, 2017; Cekanaviciute et al, 2017).

In this work, we showed that the secretome of *A. muciniphila* cultivated in mucin-free medium induces RYR-dependent ER Ca^2+^ release. This persistent Ca^2+^- mediated signals is followed by increased mitochondrial Ca^2+^ uptake in STC-1 cells, which in turn culminates with aSyn phosphorylation and aggregation. On the other hand, the secretome isolated from *A.muciniphila* grown in mucin-supplemented medium cause no intracellular alteration in the enteroendocrine cell line studied. This phenomenon leads us to the fact that in order to maintain the mammalian intestinal homeostasis with the microbiota, a key element is to minimize and regulate contact between luminal microorganisms and the intestinal epithelial cell surface. In the gut, physical separation of bacteria and the epithelium is greatly accomplished by secretion of mucus, antimicrobial proteins, and IgA into the lumen (Macpherson et al, 2000; Macpherson & Harris, 2004). Interestingly, it was previously shown that aged mice (15 – 19 months old) have an impaired mucus barrier in the colon and ileum accompanied by major changes in the fecal microbiota composition, a fact that has also been observed in humans (Elderman et al, 2017; Sovran et al, 2019). Therefore, besides changing the microbiome pattern of secretion (Fig EV1), a decrease on mucus barrier thickness leads to increased contact of gut bacteria and their secreted components with the intestinal epithelium that could therefore modulate gut cells homeostasis, especially misbalancing intracellular Ca^2+^ dynamics.

An emerging, key pathological feature in neurons affected by PD is the global dysregulation of Ca^2+^ homeostasis (Zaichick et al, 2017). Ca^2+^ handling through contact sites between the endoplasmic reticulum and mitochondria (mitochondria-associated endoplasmic reticulum membranes – MAMs) have attracted great attention in the study of cell homeostasis and dysfunction, especially in the context of neurodegenerative disorders. Emerging evidence suggests that the abnormality and dysfunction of MAMs have been involved in a number of neurodegenerative disorders including Alzheimer’s disease, amyotrophic lateral sclerosis, and Parkinson’s disease (Gautier et al, 2016; Hedskog et al, 2013; Liu & Zhu, 2017). Also, increased intracellular Ca^2+^ levels alter Ca^2+^ handling in intracellular organelles such as the endoplasmic reticulum and mitochondria which may potentiate pathological effects (Ludtmann & Abramov, 2018). Indeed, Ca^2+^ uptake into the mitochondria is a key mechanism by which cells maintain intracellular Ca^2+^ homeostasis (Dey et al, 2020; Vos et al, 2010). However, excessive mitochondrial Ca^2+^ uptake or impaired Ca^2+^ efflux results in ROS production (Luongo et al, 2017; Reynolds & Hastings, 1995) and disruption of membrane potential inducing neuronal cell death, an important indicator of several different neurological disorders including Alzheimer’s Disease (AD) and PD. Moreover, it has been suggested that impaired mitochondrial biogenesis, Ca^2+^ buffering and oxidative stress may precede the development of PD and AD pathology (Jadiya et al, 2019; Kandimalla et al, 2018; Pratico et al, 2001; Rani & Mondal, 2020). Recent studies indicate that increased intracellular free Ca^2+^ and oxidative stress synergistically augment the number of cytoplasmic αSyn-enriched aggregates *in vitro* and *in vivo* (Goodwin et al, 2013). In addition, it is also known that αSyn aggregates trigger increased mitochondrial Ca^2+^ transient and then leads to oxidative stress (Ganjam et al, 2019; Scudamore & Ciossek, 2018). Thus, an increase in oxidative stress can cause αSyn aggregation which can also induce further oxidative stress within the cell creating a positive feedback loop.

Although very clear for neuronal cells, we show here that this cascade of events is also triggered in EECs stimulated by *A. muciniphila* secretome. The outcome of this persistent Ca^2+^ mishandling is very similar to what has already been reported for neurons in PD. Additionally, since αSyn-containing EECs directly connect to enteric nerve terminals forming a neural circuit between the gut and the nervous system, influences in the gut lumen affect αSyn folding in the EECs which can then propagate to the nervous system. Hence, with this work we offer some mechanistic insights of how misfolded αSyn could be generated in EECs and then be propagated from the gut epithelium to the brain, being a possible outcome of Sporadic PD, as once hypothesized by Braak (Braak et al, 2003b). Finally, the data presented here provide further information to be considered in the design of microbial interventions for therapeutic use and sets the stage for dissection of the biochemical mechanisms underlying gut microbiome-induced- αSyn aggregation.

## MATERIALS AND METHODS

### Cell lines

STC-1 (CRL-3254) cell line was obtained from the America Type Culture Collection (ATCC). STC-1 cell was cultured in DMEM (Gibco) supplemented with 10% fetal bovine serum (FBS), 1% penicillin/streptomycin antibiotics (PSA) and incubated at 37°C with 5% CO_2_: 95% air.

Lyophilized *A. muciniphila* (DSM-22959) and *E. coli* (ATCC 25922) were purchased from DSMZ and America Type Culture Collection respectively. Strains were grown individually in pure Brain & Heart Infusion Broth (BHI) (BD, Heidelberg, Germany) or in BHI broth supplemented with 0.4% mucin (Sigma-Aldrich, MO, USA) (Derrien et al, 2004; Huo et al, 2020; Zhao et al, 2017).

### Secretome collection

Briefly, lyophilized *A. muciniphila* (DSM-22959) and *E. coli* (ATCC 25922) were grown individually in pure Brain & Heart Infusion Broth (BHI) (BD, Heidelberg, Germany) or in BHI broth supplemented with 0.4% mucin (Sigma-Aldrich, MO, USA) (Derrien et al, 2004; Huo et al, 2020; Zhao et al, 2017).

After being supplemented and sterilized, the vials containing the media were gassed with N_2_ injection system for 30 minutes, and then placed into an anaerobic chamber (Whitley DG250 Anaeorobic Workstation, Don Whitley Scientific) kindly provided by the Brazilian Bio renewables National Laboratory (LNBR, CNPEM). The bacterial inoculum was incubated for up to 72 hs at 37°C. After 12, 24, 36, 48, 60 and 72 hs of incubation, an aliquot of each of the vials was collected and evaluated under light microscopy and spectrometry for optical density (OD) measurement, using a spectrometer (Evolution™ 60S UV-Visible Spectrophotometer, Thermo Fisher Scientific Inc.) in order to monitor bacterial growth.

After 36-40 hs of incubation, conditioned (CM) and unconditioned media were collected and concentrated. Briefly, media were centrifuged for 4000 rpm at 4°C to pellet cells and the supernatant was concentrated at 4000 rpm at 4°C for 20min using Centricon® 3kD Plus-70 Centrifugal Filter Units. These conditioned (BHI CM) or unconditioned media (BHI) containing the secretome were then filtered (0.22 µm) and stored at -80°C until used.

### Mass Spectrometry

*A. muciniphila* CM and CM + 0.4% mucin were submitted to protein electrophoresis technique using 10% acrylamide-SDS page separation. After the electrophoretic run, the lanes were cut into small fragments, micro-purified, enriched, and digested using trypsin (Ishihama et. al, 2007 & Shevchenko et al. 1996, modified). Then, samples were directed to mass spectrometry (LTQ Orbitrap Velos, Thermo- Fischer). The peptides were separated with a 2-30% acetonitrile gradient in 0.1% formic acid using a PicoFrit analytical column (20 cm x 75 nm, particle size from 5 μm, New Objective) at a flow rate of 300 nL/min over 173 min. The nanoelectrospray voltage was adjusted to 2.2 kV and the source temperature was set at 275°C. All instrument methods were set in data dependent acquisition mode. The full scan MS spectra (m/z300-1600) were acquired in the Orbitrap analyzer after accumulation to a target value of 1×10^6^. The resolution in the Orbitrap was set at 60,000 and the 20 ions of the most intense peptides with states of charge ≥2 were sequentially isolated to a target value of 5,000 and fragmented into linear ion traps using low energy CID (35% normalized collision energy). The signal limit for triggering the MS/MS event was set to 1,000 counts. Dynamic exclusion was activated with an exclusion size list of 500, exclusion duration of 60 s and a repeat count of 1. An activation q = 0.25 and an activation time of 10 ms were used.

The data obtained by mass spectrometry were processed using the MaxQuant 1.3 software based on the *A. muciniphila* protein database.

### Plasmids and transfection

cDNA for the Ca^2+^ binding protein parvalbumin (PV) fused to mitochondrial targeting sequence (PV-MTS-GFP) and respective control (MTS-GFP) were kindly donated by Dr. Mateus Guerra (Yale University, USA) (Guerra et al, 2011). cDNA for the human αSyn (αSyn) was amplified and cloned between the *HindIII* and *SmaI* restriction sites of pEGFP-N2 vector. For αSyn mCherry-tagged version, the human αSyn sequence was inserted between *BamH1* and *SalI* of the pCDNA5-mCherry vector.

### Immunofluorescence

Cells were cultured onto glass-slides and fixed with 4% PFA for 20 min. Samples were blocked in PBS 1X containing 5% Normal Horse Serum and 5% bovine albumin (Sigma-Aldrich) for 1h. After washing in PBS 1X, cells were incubated with primary antibodies anti-αSyn (1:250, Abcam); anti-pser129 αSyn, (1:100; Abcam) for 2 h at room temperature, followed by PBS washes and incubation with secondary antibody (anti-mouse Alexa-488, 1:500; Thermo-Fischer) for 1 h at room temperature. Fluorescence intensity of αSyn and αSyn-p-serine-129 was quantified on at least 55 cells from 3 different experiments. Data are expressed as percentage relative to the untreated group (control). The images were obtained using a Leica SP8 confocal microscope, using a ×63 objective lens, 1.4 NA.

### Cell viability

Cellular viability was measured using the CellTiter 96® AQueous One Solution Cell Proliferation Assay (MTS) (Promega) according to the manufacturer’s protocol. STC-1 cells were treated with 1 or 10% BHI and BHI CM for 48 hs. Following this period, media was collected for CCK secretion, and cells were incubated in the presence of MTS Tetrazolium Compound for 2 hr at 37°C. Absorbance measurements (490nm) were performed using a plate reader (PerkinElmer; Waltham, MA). For analysis of secreted CCK, we used Cholecystokinin EIA Kit (Sigma-Aldrich), following manufacturer’s instructions

### Immunobloting

Cell proteins were extracted with RIPA buffer supplemented with inhibitors of proteases and phosphatases followed by centrifugation at 10,000 rpm for 10 min at 4⁰ C. Proteins were separated by SDS-PAGE in 12% Bis-Tris gels and transferred onto 0.45 µm nitrocellulose membranes (BioRad). The blots were incubated overnight at 4°C with anti-αSyn (1:100, Abcam), GATA-2 (1:1000, R and D Systems) and anti-pser129 αSyn, (1:1000; Abcam) antibodies followed by incubation with horseradish peroxidase (HRP)-conjugated secondary IgGs (anti-rabbit, 1:5000, Thermo-Fischer; anti-mouse, 1:5000, Thermo Fischer). β-actin (1:5000, Santa Cruz) or GAPDH (1:5000, Santa Cruz) were used as loading controls. Membranes were developed using the BioRad chemiluminescence detection system (Clarity Western ECL, BioRad). Chemiluminescence signals were quantified using Image J software. β -actin or GAPDH was used as a loading control.

### Cytosolic and mitochondrial calcium measurements

For cytosolic calcium measurements, cells were loaded with the Ca^2+^ indicator Fluo-4/AM (for intracellular Ca^2+^) or Rhod-2/AM (for mitochondrial Ca^2+^) (Thermo Fisher Scientific) for 15 min at 37°C, placed onto the stage of a Leica SP8 Confocal System and continuously perfused with HEPES buffer solution (142.2 mM NaCl, 5.4 mM KCl, 1.0 mM NaH_2_PO_4_, 10 mM HEPES, 5.6 mM dextrose, 0.8 mM MgSO_4_ and 1 mM CaCl_2_), unless otherwise noted. 1 or 10% BHI or BHI CM (v/v) medium with or without 0.4% mucin was used to trigger Ca^2+^ release. 40 µM Adenosine triphosphate (ATP) was used to trigger InsP3-dependent Ca^2+^ release and to evaluate mitochondrial-calcium response after 48h-treatment with the secretome. To investigate whether intracellular calcium signaling was from endoplasmic reticulum stores, cells were incubated for 30 min in HEPES Ca^2+^-free buffer containing 10µM thapsigargin prior to stimulation with the conditioned medium. To investigate IP_3_ dependence for the *A. muciniphila* conditioned media-triggered calcium response, cells were incubated with Fluo-4/AM and 2.5 µM xestospongin C for 30 min before stimulation with the conditioned medium. Dantrolene sodium salt (75 μM) was used as RYR blocker. For mitochondrial calcium measurements in PV-MTS-GFP or MTS-GFP transfected cells, cells were transfected 48 hs before the experiment using FUGENE HD (Promega) according to the manufacturer’s instructions. Data are expressed as fluorescence/baseline fluorescence × 100% of the average values of samples from 3-6 biological replicates (>20 cells/replicate). The images were obtained using a Leica SP8 confocal microscope, using a ×63 objective lens, 1.4 NA, excitation at 488 nm and emission at 505-525 nm for both dyes.

### Detection of reactive oxygen species (ROS)

To evaluate the formation of reactive oxygen species (ROS), STC-1 cells were previously seeded on glass slides were treated with 5 µM DHE (Thermo-Fischer) for 30 min in HEPES buffer solution (142.2 mM NaCl, 5.4 mM KCl, 1.0 mM NaH_2_PO_4_, 10 mM HEPES, 5.6 mM dextrose, 0.8 mM MgSO_4_ and 1 mM CaCl_2_). The glass slides were transferred to a perfusion chamber attached to the confocal microscope. The cells were stimulated with 1 or 10% (v/v) *A. muciniphila* conditioned or unconditioned media. As a positive control of ROS formation, 100 µM H_2_O_2_ was used.

For each stimulus, the emission of fluorescence in response to the general indicator of oxidative stress was monitored in individualized cells during stimulation with the conditioned and unconditioned media. Data are expressed as fluorescence/baseline fluorescence × 100% of the average values of samples from 3-6 biological replicates (>20 cells/replicate). The images were obtained using a Leica SP8 confocal microscope, using a ×63 objective lens, 1.4 NA, excitation at 488-518 nm and emission at 606 nm.

### Evaluation of mitochondrial membrane potential

STC-1 cells were treated for 48 hs with 1 or 10% (v/v) *A. muciniphila* conditioned or unconditioned media and then incubated 500 nM of MitoTracker Red CMXRos (Thermo-Fischer) for 30 min. Cells were washed 2x with PBS1x and then fixed in 4% paraformaldehyde at room temperature, for 20 min, washed with PBS and in sequence, were mounted in Vectashield. At least 55 cells from 3 different experiments were analyzed. Data are expressed as percentage relative to the untreated group (control). The images were obtained using a Leica SP8 confocal microscope, using a ×63 objective lens, 1.4 NA, excitation at 579 nm and emission at 599 nm. Finally, the intensity of fluorescence of the dye was quantified

### Statistical analysis

The statistical relevance of the bar graphs was obtained by calculating the P-value using the paired two-tailed Student’s t-test. The bar graphs showed in the Figures are presented as mean ± SEM. For the imaging experiments, at least 55 independent cells for each condition were analyzed. Comparison of multiple groups was performed by one-way analysis of variance with Bonferroni post-tests. All column graphs, plots and statistical analyses were done using GraphPad Prism version 6 software.

## ACKNOWLEDGEMENTS

We thank Dr. Kleber Gomes Franchini (LNBio-CNPEM, Brazil) for his contribution in the initial phase of this project and for carefully reading the manuscript. We also thank Dr. Hernandes Faustino Carvalho (Unicamp, Brazil) for valuable comments on the manuscript. The authors would like to acknowledge support of the Brazilian Biorenewables National Laboratory (LNBR – CNPEM) for providing access to its facilities. This work was supported by FAPESP (2018/20014-0; and 2019/24511-0).

## AUTHOR CONTRIBUTIONS

Conceptualisation, MCF and DPAN; Formal Analysis, DPAN, BPG, and MCF; Investigation DPAN, BPB, JVPG, KT, PV, CCCT; Resources, CCCT and MCF; Writing, DPAN, CGB and MCF; Funding Acquisition, MCF.

## DATA AND CODE AVAILABILITY

All developed expression plasmids produced in this study can be made available upon request to the correspondent author. This study did not generate any unique datasets or code. All data supporting the findings of this study are available within the paper and are available from the corresponding author upon request.

## SUPPORTING INFORMATION

Expanded View Figs 1 – 6.

**Expanded View Fig 1. Growth and proteomics curve of *A. muciniphila* secretome cultivated in the presence or absence of 0.4% mucin.**

**A.** Growth curve as a function of culture media supplementation. Error bars indicate the ± SEM of three individual bacterial culture for each condition.

**B.** Venn diagram showing the common and unique expressed proteins between the 0.4% mucin and mucin-free culture condition identified by mass spectrometry.

Data information: Growth curves and mass spectrometry data were obtained from at least 6 vials for each culture condition. Data in (A) represent the mean ± SEM of at least vials of each culture condition.

**Expanded View Fig 2. 0.4% mucin-supplemented *Akkermansia muciniphila* conditioned medium induces discrete intracellular calcium signals and does not influence total levels of α-synuclein in STC-1 cells.**

**A.** Confocal microscopy imaging of STC-1 cells incubated with Fluo-4/AM (6μM) and stimulated with 1 or 10% 0.4% mucin-supplemented *A. muciniphila* conditioned media (BHI CM + 0.4% mucin) (scale bar: 10 μm).

**B.** Representative time-course of total Ca^2+^ signal. The arrow indicates the time when culture medium was applied.

**C.** Quantification of the peak fluorescence following stimulation with 1 or 10% conditioned (BHI CM) and unconditioned (BHI) media. * P < 0.05; Ns, not significant by two-way Student’s *t-test*.

**D.** αSyn staining (green) in STC-1 cells after 48 hs incubation with 1-10% conditioned (BHI CM) or unconditioned mucin-supplemented media (BHI) demonstrating no differential expression of the protein. Nuclei were stained with DAPI (blue) and immunofluorescence control is shown as NSB (non-specific binding control). (scale bar: 10 µm).

**E.** Quantification of αSyn fluorescence intensity in images shown in (D).

**F.** Immunoblots (upper image) of total cell lysates showing expression of αSyn after 48 hours-incubation with 1-10% conditioned (BHI CM) or unconditioned mucin- supplemented media (BHI). Graphic representation of densitometric analysis showing expression of αSyn in 1-10% BHI and BHI CM mucin-supplemented condition.

Data information: Data in (B) represent a representative tracing recorded from one individual STC-1 cell of each group. Data in (C and E-G) represent the mean ± SEM of three independent experiments. Whereas at least 55 individual cells were analyzed for calcium transient experiments and immunofluorescence, densitometric analysis of western blot are derived from triplicates of three different experiments.

**Expanded View Fig 3. Mucin-free or mucin-supplemented media do not induce intracellular calcium signals in STC-1 cells.**

**A.** Confocal microscopy imaging of STC-1 cells incubated with Fluo-4/AM (6 μM) and stimulated with 1 or 10% unconditioned mucin-free media (BHI) (scale bar: 10 μm).

**B.** Representative time-course of total Ca^2+^ signal. Arrow indicates the moment when culture medium was applied.

**C.** Quantification of the peak fluorescence following stimulation with 1 or 10% unconditioned mucin-free media (BHI). Ns, not significant by two-way Student’s *t-test*.

**D.** Confocal microscopy imaging of STC-1 cells incubated with Fluo-4/AM (6μM) and stimulated with 1 or 10% unconditioned 0.4% mucin-supplemented media (BHI + 0.4% mucin) (scale bar: 10 μm).

**E.** Representative time-course of total Ca^2+^ signal. The arrow indicates the time when culture medium was applied.

**F.** Quantification of the peak fluorescence following stimulation with 1 or 10% unconditioned 0.4% mucin-supplemented media (BHI + 0.4% mucin). Ns, not significant by two-way Student’s *t-test*.

Data information: Data in (B and E) represent a representative tracing recorded from one individual STC-1 cell of each group. Data in (C and F) represent the mean ± SEM of three independent experiments in which at least 55 individual cells were analyzed.

**Expanded View Fig 4. Incubation of STC-1 cells with *Akkermansia muciniphila* conditioned or unconditioned media does not alter cell viability.**

**A.** MTS assay expressed as percentage of control showing no reduction of STC-1 cell viability after treatment. 1% Triton-X 100 was used as positive control. * P < 0.001 by one-way ANOVA.

**B.** Cholecystokinin (CCK) secretion measured in extracellular medium by ELISA after 48hrs incubation with 1-10% *A. muciniphila* unconditioned (BHI) or conditioned (BHI CM) media.

Data information: Values for toxicity and ELISA experiments are derived from triplicates of three different experiments.

**Expanded View Fig 5. Although *Escherichia coli* conditioned medium induces intracellular calcium signals, it does not influence total levels of α-synuclein in STC-1 cells.**

**A-B.** Confocal microscopy imaging of STC-1 cells incubated with Fluo-4/AM (6 μM) and stimulated with 1 **(A)** or 10% **(B)** *E. coli* conditioned (BHI CM) media (scale bar: 10 μm). * P < 0.05 by two-way Student’s *t-test*.

**C-D.** Representative time-course of total Ca^2+^ signal. The arrow indicates the time when *E. coli* conditioned (BHI CM) (**C)** or unconditioned (BHI) **(D)** culture medium was applied.

**E.** Quantification of the peak fluorescence following stimulation with 1 or 10% *E. coli* and *A. muciniphila* conditioned (BHI CM) and unconditioned (BHI) media.

**F.** αSyn staining (green) in STC-1 cells after 48 hs incubation with 1-10% *E. coli* conditioned (BHI CM) media demonstrating no alteration in the protein expression. Nuclei were stained with DAPI (blue) (scale bar: 10 µm)

**G.** Quantification of αSyn fluorescence intensity in images shown in (D).

**H.** Immunoblots (upper image) of total cell lysates showing expression of αSyn after 48 hours-incubation with 1-10% *E. coli* conditioned (BHI CM) or unconditioned media (BHI). Graphic representation of densitometric analysis showing expression of αSyn in 1/10% BHI CM or BHI media.

Data information: Data in (C and D) represent a representative tracing recorded from one individual STC-1 cell of each group. Data in (E) represent the mean ± SEM of three independent experiments. Whereas at least 55 individual cells were analyzed for calcium transient experiments and immunofluorescence (G), densitometric analysis of western blot are derived from triplicates of three different experiments (H).

**Expanded View Fig 6. *A. muciniphila* conditioned media promotes Ca^2+^ increase with greater intensity in the cytoplasm when compared to the nuclear region.**

**A.** Line scan of Ca^2+^ signal in STC-1 cells loaded with Fluo-4/AM. It is observed that *A. muciniphila* conditioned (BHI CM) media promotes Ca^2+^ increase with greater intensity in the cytoplasm when compared to the nuclear region.

**B.** Line tracing of the fluorescence Ca^2+^ intensity throughout a representative cell (blue line in image A) shows increased fluorescence signal in the cytoplasm after stimulus (black line when compared to gray line).

Data information: Data in (B) represent a representative tracing recorded from one individual STC-1 cell showed in A.

## REFERENCES

1. Abbott RD, Petrovitch H, White LR, Masaki KH, Tanner CM, Curb JD, Grandinetti A, Blanchette PL, Popper JS, Ross GW (2001) Frequency of bowel movements and the future risk of Parkinson’s disease. Neurology 57: 456–462

2. Adams-Carr KL, Bestwick JP, Shribman S, Lees A, Schrag A, Noyce AJ (2016) Constipation preceding Parkinson’s disease: a systematic review and meta-analysis. Journal of neurology, neurosurgery, and psychiatry 87: 710–716

3. Alvarenga EC, Fonseca MC, Carvalho CC, Florentino RM, Franca A, Matias E, Guimaraes PB, Batista C, Freire V, Carmona AK, Pesquero JB, de Paula AM, Foureaux G, Leite MF (2016) Angiotensin Converting Enzyme Regulates Cell Proliferation and Migration. PloS one 11: e0165371

4. Baldini F, Hertel J, Sandt E, Thinnes CC, Neuberger-Castillo L, Pavelka L, Betsou F, Kruger R, Thiele I, Consortium N-P (2020) Parkinson’s disease-associated alterations of the gut microbiome predict disease-relevant changes in metabolic functions. BMC biology 18: 62

5. Bedarf JR, Hildebrand F, Coelho LP, Sunagawa S, Bahram M, Goeser F, Bork P, Wullner U (2017) Functional implications of microbial and viral gut metagenome changes in early stage L-DOPA-naive Parkinson’s disease patients. Genome medicine 9: 39

6. Bekris LM, Mata IF, Zabetian CP (2010) The genetics of Parkinson disease. Journal of geriatric psychiatry and neurology 23: 228–242

7. Berer K, Gerdes LA, Cekanaviciute E, Jia X, Xiao L, Xia Z, Liu C, Klotz L, Stauffer U, Baranzini SE, Kumpfel T, Hohlfeld R, Krishnamoorthy G, Wekerle H (2017) Gut microbiota from multiple sclerosis patients enables spontaneous autoimmune encephalomyelitis in mice. Proceedings of the National Academy of Sciences of the United States of America 114: 10719–10724

8. Bernheimer H, Birkmayer W, Hornykiewicz O, Jellinger K, Seitelberger F (1973) Brain dopamine and the syndromes of Parkinson and Huntington. Clinical, morphological and neurochemical correlations. Journal of the neurological sciences 20: 415–455

9. Bilodeau ML, Boulineau T, Greulich JD, Hullinger RL, Andrisani OM (2001) Differential expression of sympathoadrenal lineage-determining genes and phenotypic markers in cultured primary neural crest cells. In vitro cellular & developmental biology Animal 37: 185–192

10. Bohorquez DV, Shahid RA, Erdmann A, Kreger AM, Wang Y, Calakos N, Wang F, Liddle RA (2015) Neuroepithelial circuit formed by innervation of sensory enteroendocrine cells. The Journal of clinical investigation 125: 782–786

11. Braak H, Del Tredici K (2009) Neuroanatomy and pathology of sporadic Parkinson’s disease. Advances in anatomy, embryology, and cell biology 201: 1–119

12. Braak H, Del Tredici K, Rub U, de Vos RA, Jansen Steur EN, Braak E (2003a) Staging of brain pathology related to sporadic Parkinson’s disease. Neurobiology of aging 24: 197–211

13. Braak H, Rub U, Gai WP, Del Tredici K (2003b) Idiopathic Parkinson’s disease: possible routes by which vulnerable neuronal types may be subject to neuroinvasion by an unknown pathogen. Journal of neural transmission 110: 517–536

14. Calvo-Rodriguez M, Hou SS, Snyder AC, Kharitonova EK, Russ AN, Das S, Fan Z, Muzikansky A, Garcia-Alloza M, Serrano-Pozo A, Hudry E, Bacskai BJ (2020) Increased mitochondrial calcium levels associated with neuronal death in a mouse model of Alzheimer’s disease. Nature communications 11: 2146

15. Carafoli E, Krebs J (2016) Why Calcium? How Calcium Became the Best Communicator. The Journal of biological chemistry 291: 20849–20857

16. Cekanaviciute E, Yoo BB, Runia TF, Debelius JW, Singh S, Nelson CA, Kanner R, Bencosme Y, Lee YK, Hauser SL, Crabtree-Hartman E, Sand IK, Gacias M, Zhu Y, Casaccia P, Cree BAC, Knight R, Mazmanian SK, Baranzini SE (2017) Gut bacteria from multiple sclerosis patients modulate human T cells and exacerbate symptoms in mouse models. Proceedings of the National Academy of Sciences of the United States of America 114: 10713–10718

17. Cersosimo MG, Raina GB, Pecci C, Pellene A, Calandra CR, Gutierrez C, Micheli FE, Benarroch EE (2013) Gastrointestinal manifestations in Parkinson’s disease: prevalence and occurrence before motor symptoms. Journal of neurology 260: 1332–1338

18. Chandra R, Hiniker A, Kuo YM, Nussbaum RL, Liddle RA (2017) alpha-Synuclein in gut endocrine cells and its implications for Parkinson’s disease. JCI insight 2

19. Chaudhuri KR, Healy DG, Schapira AH, National Institute for Clinical E (2006) Non-motor symptoms of Parkinson’s disease: diagnosis and management. The Lancet Neurology 5: 235–245

20. Davie CA (2008) A review of Parkinson’s disease. British medical bulletin 86: 109–127

21. de Lau LM, Breteler MM (2006) Epidemiology of Parkinson’s disease. The Lancet Neurology 5: 525–535

22. Derrien M, Belzer C, de Vos WM (2017) Akkermansia muciniphila and its role in regulating host functions. Microbial pathogenesis 106: 171–181

23. Derrien M, Vaughan EE, Plugge CM, de Vos WM (2004) Akkermansia muciniphila gen. nov., sp. nov., a human intestinal mucin-degrading bacterium. International journal of systematic and evolutionary microbiology 54: 1469–1476

24. Desai MS, Seekatz AM, Koropatkin NM, Kamada N, Hickey CA, Wolter M, Pudlo NA, Kitamoto S, Terrapon N, Muller A, Young VB, Henrissat B, Wilmes P, Stappenbeck TS, Nunez G, Martens EC (2016) A Dietary Fiber-Deprived Gut Microbiota Degrades the Colonic Mucus Barrier and Enhances Pathogen Susceptibility. Cell 167: 1339–1353 e1321

25. Dey K, Bazala MA, Kuznicki J (2020) Targeting mitochondrial calcium pathways as a potential treatment against Parkinson’s disease. Cell calcium 89: 102216

26. Di Lisa F, Bernardi P (2009) A CaPful of mechanisms regulating the mitochondrial permeability transition. Journal of molecular and cellular cardiology 46: 775–780

27. Divecha N, Banfic H, Irvine RF (1991) The polyphosphoinositide cycle exists in the nuclei of Swiss 3T3 cells under the control of a receptor (for IGF-I) in the plasma membrane, and stimulation of the cycle increases nuclear diacylglycerol and apparently induces translocation of protein kinase C to the nucleus. The EMBO journal 10: 3207–3214

28. Elderman M, Sovran B, Hugenholtz F, Graversen K, Huijskes M, Houtsma E, Belzer C, Boekschoten M, de Vos P, Dekker J, Wells J, Faas M (2017) The effect of age on the intestinal mucus thickness, microbiota composition and immunity in relation to sex in mice. PloS one 12: e0184274

29. Fonseca MC, Franca A, Florentino RM, Fonseca RC, Lima Filho ACM, Vidigal PTV, Oliveira AG, Dubuquoy L, Nathanson MH, Leite MF (2018) Cholesterol-enriched membrane microdomains are needed for insulin signaling and proliferation in hepatic cells. American journal of physiology Gastrointestinal and liver physiology 315: G80–G94

30. Gafni J, Munsch JA, Lam TH, Catlin MC, Costa LG, Molinski TF, Pessah IN (1997) Xestospongins: potent membrane permeable blockers of the inositol 1,4,5-trisphosphate receptor. Neuron 19: 723–733

31. Gagic D, Ciric M, Wen WX, Ng F, Rakonjac J (2016) Exploring the Secretomes of Microbes and Microbial Communities Using Filamentous Phage Display. Frontiers in microbiology 7: 429

32. Ganjam GK, Bolte K, Matschke LA, Neitemeier S, Dolga AM, Hollerhage M, Hoglinger GU, Adamczyk A, Decher N, Oertel WH, Culmsee C (2019) Mitochondrial damage by alpha-synuclein causes cell death in human dopaminergic neurons. Cell death & disease 10: 865

33. Gautier CA, Erpapazoglou Z, Mouton-Liger F, Muriel MP, Cormier F, Bigou S, Duffaure S, Girard M, Foret B, Iannielli A, Broccoli V, Dalle C, Bohl D, Michel PP, Corvol JC, Brice A, Corti O (2016) The endoplasmic reticulum-mitochondria interface is perturbed in PARK2 knockout mice and patients with PARK2 mutations. Human molecular genetics 25: 2972–2984

34. Goedert M, Spillantini MG, Del Tredici K, Braak H (2013) 100 years of Lewy pathology. Nature reviews Neurology 9: 13–24

35. Goodwin J, Nath S, Engelborghs Y, Pountney DL (2013) Raised calcium and oxidative stress cooperatively promote alpha-synuclein aggregate formation. Neurochemistry international 62: 703–711

36. Grander C, Adolph TE, Wieser V, Lowe P, Wrzosek L, Gyongyosi B, Ward DV, Grabherr F, Gerner RR, Pfister A, Enrich B, Ciocan D, Macheiner S, Mayr L, Drach M, Moser P, Moschen AR, Perlemuter G, Szabo G, Cassard AM, Tilg H (2018) Recovery of ethanol-induced Akkermansia muciniphila depletion ameliorates alcoholic liver disease. Gut 67: 891–901

37. Greenamyre JT, Hastings TG (2004) Biomedicine. Parkinson’s--divergent causes, convergent mechanisms. Science 304: 1120–1122

38. Greenland JC, Williams-Gray CH, Barker RA (2019) The clinical heterogeneity of Parkinson’s disease and its therapeutic implications. The European journal of neuroscience 49: 328–338

39. Guerra MT, Fonseca EA, Melo FM, Andrade VA, Aguiar CJ, Andrade LM, Pinheiro AC, Casteluber MC, Resende RR, Pinto MC, Fernandes SO, Cardoso VN, Souza- Fagundes EM, Menezes GB, de Paula AM, Nathanson MH, Leite Mde F (2011) Mitochondrial calcium regulates rat liver regeneration through the modulation of apoptosis. Hepatology 54: 296–306

40. Guimaraes E, Machado R, Fonseca MC, Franca A, Carvalho C, Araujo ESAC, Almeida B, Cassini P, Hissa B, Drumond L, Goncalves C, Fernandes G, De Brot M, Moraes M, Barcelos L, Ortega JM, Oliveira A, Leite MF (2017) Inositol 1, 4, 5-trisphosphate- dependent nuclear calcium signals regulate angiogenesis and cell motility in triple negative breast cancer. PloS one 12: e0175041

41. Gunter TE, Buntinas L, Sparagna GC, Gunter KK (1998) The Ca2+ transport mechanisms of mitochondria and Ca2+ uptake from physiological-type Ca2+ transients. Biochimica et biophysica acta 1366: 5–15

42. Guzman JN, Sanchez-Padilla J, Wokosin D, Kondapalli J, Ilijic E, Schumacker PT, Surmeier DJ (2010) Oxidant stress evoked by pacemaking in dopaminergic neurons is attenuated by DJ-1. Nature 468: 696–700

43. Hainaut K, Desmedt JE (1974) Effect of dantrolene sodium on calcium movements in single muscle fibres. Nature 252: 728–730

44. Hand KV, Bruen CM, O’Halloran F, Panwar H, Calderwood D, Giblin L, Green BD (2013) Examining acute and chronic effects of short- and long-chain fatty acids on peptide YY (PYY) gene expression, cellular storage and secretion in STC-1 cells. European journal of nutrition 52: 1303–1313

45. Hand KV, Giblin L, Green BD (2012) Hormone profiling in a novel enteroendocrine cell line pGIP/neo: STC-1. Metabolism: clinical and experimental 61: 1683–1686

46. Hawkes CH, Del Tredici K, Braak H (2010) A timeline for Parkinson’s disease. Parkinsonism & related disorders 16: 79–84

47. Hedskog L, Pinho CM, Filadi R, Ronnback A, Hertwig L, Wiehager B, Larssen P, Gellhaar S, Sandebring A, Westerlund M, Graff C, Winblad B, Galter D, Behbahani H, Pizzo P, Glaser E, Ankarcrona M (2013) Modulation of the endoplasmic reticulum-mitochondria interface in Alzheimer’s disease and related models. Proceedings of the National Academy of Sciences of the United States of America 110: 7916–7921

48. Heintz-Buschart A, Pandey U, Wicke T, Sixel-Doring F, Janzen A, Sittig-Wiegand E, Trenkwalder C, Oertel WH, Mollenhauer B, Wilmes P (2018) The nasal and gut microbiome in Parkinson’s disease and idiopathic rapid eye movement sleep behavior disorder. Movement disorders : official journal of the Movement Disorder Society 33: 88–98

49. Hijaz BA, Volpicelli-Daley LA (2020) Initiation and propagation of alpha-synuclein aggregation in the nervous system. Molecular neurodegeneration 15: 19

50. Hill-Burns EM, Debelius JW, Morton JT, Wissemann WT, Lewis MR, Wallen ZD, Peddada SD, Factor SA, Molho E, Zabetian CP, Knight R, Payami H (2017) Parkinson’s disease and Parkinson’s disease medications have distinct signatures of the gut microbiome. Movement disorders : official journal of the Movement Disorder Society 32: 739–749

51. Hill BG, Benavides GA, Lancaster JR, Jr., Ballinger S, Dell’Italia L, Jianhua Z, Darley-Usmar VM (2012) Integration of cellular bioenergetics with mitochondrial quality control and autophagy. Biological chemistry 393: 1485–1512

52. Huo Y, Lu X, Wang X, Wang X, Chen L, Guo H, Zhang M, Li Y (2020) Bifidobacterium animalis subsp. lactis A6 Alleviates Obesity Associated with Promoting Mitochondrial Biogenesis and Function of Adipose Tissue in Mice. Molecules 25

53. Jacotot E, Ferri KF, Kroemer G (2000a) Apoptosis and cell cycle: distinct checkpoints with overlapping upstream control. Pathologie-biologie 48: 271–279

54. Jacotot E, Ravagnan L, Loeffler M, Ferri KF, Vieira HL, Zamzami N, Costantini P, Druillennec S, Hoebeke J, Briand JP, Irinopoulou T, Daugas E, Susin SA, Cointe D, Xie ZH, Reed JC, Roques BP, Kroemer G (2000b) The HIV-1 viral protein R induces apoptosis via a direct effect on the mitochondrial permeability transition pore. The Journal of experimental medicine 191: 33–46

55. Jadiya P, Kolmetzky DW, Tomar D, Di Meco A, Lombardi AA, Lambert JP, Luongo TS, Ludtmann MH, Pratico D, Elrod JW (2019) Impaired mitochondrial calcium efflux contributes to disease progression in models of Alzheimer’s disease. Nature communications 10: 3885

56. Jost WH (2010) Gastrointestinal dysfunction in Parkinson’s Disease. Journal of the neurological sciences 289: 69–73

57. Kandimalla R, Manczak M, Yin X, Wang R, Reddy PH (2018) Hippocampal phosphorylated tau induced cognitive decline, dendritic spine loss and mitochondrial abnormalities in a mouse model of Alzheimer’s disease. Human molecular genetics 27: 30–40

58. Karampetsou M, Ardah MT, Semitekolou M, Polissidis A, Samiotaki M, Kalomoiri M, Majbour N, Xanthou G, El-Agnaf OMA, Vekrellis K (2017) Phosphorylated exogenous alpha-synuclein fibrils exacerbate pathology and induce neuronal dysfunction in mice. Scientific reports 7: 16533

59. Keshavarzian A, Green SJ, Engen PA, Voigt RM, Naqib A, Forsyth CB, Mutlu E, Shannon KM (2015) Colonic bacterial composition in Parkinson’s disease. Movement disorders : official journal of the Movement Disorder Society 30: 1351–1360

60. Kim S, Kwon SH, Kam TI, Panicker N, Karuppagounder SS, Lee S, Lee JH, Kim WR, Kook M, Foss CA, Shen C, Lee H, Kulkarni S, Pasricha PJ, Lee G, Pomper MG, Dawson VL, Dawson TM, Ko HS (2019) Transneuronal Propagation of Pathologic alpha-Synuclein from the Gut to the Brain Models Parkinson’s Disease. Neuron 103: 627–641 e627

61. Klingelhoefer L, Reichmann H (2015) Pathogenesis of Parkinson disease--the gut-brain axis and environmental factors. Nature reviews Neurology 11: 625–636

62. Li W, Wu X, Hu X, Wang T, Liang S, Duan Y, Jin F, Qin B (2017) Structural changes of gut microbiota in Parkinson’s disease and its correlation with clinical features. Science China Life sciences 60: 1223–1233

63. Liddle RA (2018) Parkinson’s disease from the gut. Brain research 1693: 201–206

64. Lin CH, Chen CC, Chiang HL, Liou JM, Chang CM, Lu TP, Chuang EY, Tai YC, Cheng C, Lin HY, Wu MS (2019) Altered gut microbiota and inflammatory cytokine responses in patients with Parkinson’s disease. Journal of neuroinflammation 16: 129

65. Liu Y, Zhu X (2017) Endoplasmic reticulum-mitochondria tethering in neurodegenerative diseases. Translational neurodegeneration 6: 21

66. Ludtmann MHR, Abramov AY (2018) Mitochondrial calcium imbalance in Parkinson’s disease. Neuroscience letters 663: 86–90

67. Luongo TS, Lambert JP, Gross P, Nwokedi M, Lombardi AA, Shanmughapriya S, Carpenter AC, Kolmetzky D, Gao E, van Berlo JH, Tsai EJ, Molkentin JD, Chen X, Madesh M, Houser SR, Elrod JW (2017) The mitochondrial Na(+)/Ca(2+) exchanger is essential for Ca(2+) homeostasis and viability. Nature 545: 93–97

68. Macpherson AJ, Gatto D, Sainsbury E, Harriman GR, Hengartner H, Zinkernagel RM (2000) A primitive T cell-independent mechanism of intestinal mucosal IgA responses to commensal bacteria. Science 288: 2222–2226

69. Macpherson AJ, Harris NL (2004) Interactions between commensal intestinal bacteria and the immune system. Nature reviews Immunology 4: 478–485

70. Martinez-Martin P, Rodriguez-Blazquez C, Kurtis MM, Chaudhuri KR, Group NV (2011) The impact of non-motor symptoms on health-related quality of life of patients with Parkinson’s disease. Movement disorders : official journal of the Movement Disorder Society 26: 399–406

71. McCarthy T, Green BD, Calderwood D, Gillespie A, Cryan JF, Giblin L (2015) STC-1 Cells. In The Impact of Food Bioactives on Health: in vitro and ex vivo models, Verhoeckx K, Cotter P, Lopez-Exposito I, Kleiveland C, Lea T, Mackie A, Requena T, Swiatecka D, Wichers H (eds), pp 211–220. Cham (CH)

72. Mertsalmi TH, Aho VTE, Pereira PAB, Paulin L, Pekkonen E, Auvinen P, Scheperjans F (2017) More than constipation - bowel symptoms in Parkinson’s disease and their connection to gut microbiota. European journal of neurology 24: 1375–1383

73. Morgan KG, Bryant SH (1977) The mechanism of action of dantrolene sodium. The Journal of pharmacology and experimental therapeutics 201: 138–147

74. Mun JK, Youn J, Cho JW, Oh ES, Kim JS, Park S, Jang W, Park JS, Koh SB, Lee JH, Park HK, Kim HJ, Jeon BS, Shin HW, Choi SA, Kim SJ, Choi SM, Park JY, Kim JY, Chung SJ, Lee CS, Ahn TB, Kim WC, Kim HS, Cheon SM, Kim JW, Kim HT, Lee JY, Kim JS, Kim EJ, Kim JM, Lee KS, Kim JS, Kim MJ, Baik JS, Park KJ, Kim HJ, Park MY, Kang JH, Song SK, Kim YD, Yun JY, Lee HW, Song IU, Sohn YH, Lee PH, Park JH, Oh HG, Park KW, Kwon DY (2016) Weight Change Is a Characteristic Non-Motor Symptom in Drug-Naive Parkinson’s Disease Patients with Non-Tremor Dominant Subtype: A Nation-Wide Observational Study. PloS one 11: e0162254

75. Musgrove RE, Helwig M, Bae EJ, Aboutalebi H, Lee SJ, Ulusoy A, Di Monte DA (2019) Oxidative stress in vagal neurons promotes parkinsonian pathology and intercellular alpha-synuclein transfer. The Journal of clinical investigation 129: 3738–3753

76. Nardelli J, Thiesson D, Fujiwara Y, Tsai FY, Orkin SH (1999) Expression and genetic interaction of transcription factors GATA-2 and GATA-3 during development of the mouse central nervous system. Developmental biology 210: 305–321

77. Nath S, Goodwin J, Engelborghs Y, Pountney DL (2011) Raised calcium promotes alpha-synuclein aggregate formation. Molecular and cellular neurosciences 46: 516–526

78. Nishiwaki H, Ito M, Ishida T, Hamaguchi T, Maeda T, Kashihara K, Tsuboi Y, Ueyama J, Shimamura T, Mori H, Kurokawa K, Katsuno M, Hirayama M, Ohno K (2020) Meta- Analysis of Gut Dysbiosis in Parkinson’s Disease. Movement disorders : official journal of the Movement Disorder Society 35: 1626–1635

79. Perfeito R, Lazaro DF, Outeiro TF, Rego AC (2014) Linking alpha-synuclein phosphorylation to reactive oxygen species formation and mitochondrial dysfunction in SH-SY5Y cells. Molecular and cellular neurosciences 62: 51–59

80. Pratico D, Uryu K, Leight S, Trojanoswki JQ, Lee VM (2001) Increased lipid peroxidation precedes amyloid plaque formation in an animal model of Alzheimer amyloidosis. The Journal of neuroscience : the official journal of the Society for Neuroscience 21: 4183–4187

81. Rani L, Mondal AC (2020) Emerging concepts of mitochondrial dysfunction in Parkinson’s disease progression: Pathogenic and therapeutic implications. Mitochondrion 50: 25–34

82. Reynolds IJ, Hastings TG (1995) Glutamate induces the production of reactive oxygen species in cultured forebrain neurons following NMDA receptor activation. The Journal of neuroscience : the official journal of the Society for Neuroscience 15: 3318–3327

83. Scheperjans F, Aho V, Pereira PA, Koskinen K, Paulin L, Pekkonen E, Haapaniemi E, Kaakkola S, Eerola-Rautio J, Pohja M, Kinnunen E, Murros K, Auvinen P (2015) Gut microbiota are related to Parkinson’s disease and clinical phenotype. Movement disorders : official journal of the Movement Disorder Society 30: 350–358

84. Scherzer CR, Grass JA, Liao Z, Pepivani I, Zheng B, Eklund AC, Ney PA, Ng J, McGoldrick M, Mollenhauer B, Bresnick EH, Schlossmacher MG (2008) GATA transcription factors directly regulate the Parkinson’s disease-linked gene alpha-synuclein. Proceedings of the National Academy of Sciences of the United States of America 105: 10907–10912

85. Schneeberger M, Everard A, Gomez-Valades AG, Matamoros S, Ramirez S, Delzenne NM, Gomis R, Claret M, Cani PD (2015) Akkermansia muciniphila inversely correlates with the onset of inflammation, altered adipose tissue metabolism and metabolic disorders during obesity in mice. Scientific reports 5: 16643

86. Scudamore O, Ciossek T (2018) Increased Oxidative Stress Exacerbates alpha-Synuclein Aggregation In Vivo. Journal of neuropathology and experimental neurology 77: 443–453

87. Sovran B, Hugenholtz F, Elderman M, Van Beek AA, Graversen K, Huijskes M, Boekschoten MV, Savelkoul HFJ, De Vos P, Dekker J, Wells JM (2019) Age-associated Impairment of the Mucus Barrier Function is Associated with Profound Changes in Microbiota and Immunity. Scientific reports 9: 1437

88. Spillantini MG, Schmidt ML, Lee VM, Trojanowski JQ, Jakes R, Goedert M (1997) Alpha-synuclein in Lewy bodies. Nature 388: 839–840

89. Stefanis L (2012) alpha-Synuclein in Parkinson’s disease. Cold Spring Harbor perspectives in medicine 2: a009399

90. Sundaresan S, Shahid R, Riehl TE, Chandra R, Nassir F, Stenson WF, Liddle RA, Abumrad NA (2013) CD36-dependent signaling mediates fatty acid-induced gut release of secretin and cholecystokinin. FASEB journal : official publication of the Federation of American Societies for Experimental Biology 27: 1191–1202

91. Surmeier DJ, Schumacker PT (2013) Calcium, bioenergetics, and neuronal vulnerability in Parkinson’s disease. The Journal of biological chemistry 288: 10736–10741

92. Tenreiro S, Eckermann K, Outeiro TF (2014) Protein phosphorylation in neurodegeneration: friend or foe? Frontiers in molecular neuroscience 7: 42

93. Tinel H, Cancela JM, Mogami H, Gerasimenko JV, Gerasimenko OV, Tepikin AV, Petersen OH (1999) Active mitochondria surrounding the pancreatic acinar granule region prevent spreading of inositol trisphosphate-evoked local cytosolic Ca(2+) signals. The EMBO journal 18: 4999–5008

94. Tjalsma H, Bolhuis A, Jongbloed JD, Bron S, van Dijl JM (2000) Signal peptide- dependent protein transport in Bacillus subtilis: a genome-based survey of the secretome. Microbiology and molecular biology reviews : MMBR 64: 515–547

95. Tsarovina K, Pattyn A, Stubbusch J, Muller F, van der Wees J, Schneider C, Brunet JF, Rohrer H (2004) Essential role of Gata transcription factors in sympathetic neuron development. Development 131: 4775–4786

96. Unger MM, Spiegel J, Dillmann KU, Grundmann D, Philippeit H, Burmann J, Fassbender K, Schwiertz A, Schafer KH (2016) Short chain fatty acids and gut microbiota differ between patients with Parkinson’s disease and age-matched controls. Parkinsonism & related disorders 32: 66–72

97. Vidal-Martinez G, Chin B, Camarillo C, Herrera GV, Yang B, Sarosiek I, Perez RG (2020) A Pilot Microbiota Study in Parkinson’s Disease Patients versus Control Subjects, and Effects of FTY720 and FTY720-Mitoxy Therapies in Parkinsonian and Multiple System Atrophy Mouse Models. Journal of Parkinson’s disease 10: 185–192

98. Vos M, Lauwers E, Verstreken P (2010) Synaptic mitochondria in synaptic transmission and organization of vesicle pools in health and disease. Frontiers in synaptic neuroscience 2: 139

99. Wakabayashi K, Mori F, Tanji K, Orimo S, Takahashi H (2010) Involvement of the peripheral nervous system in synucleinopathies, tauopathies and other neurodegenerative proteinopathies of the brain. Acta neuropathologica 120: 1–12

100. Wang Y, Prpic V, Green GM, Reeve JR, Jr., Liddle RA (2002) Luminal CCK-releasing factor stimulates CCK release from human intestinal endocrine and STC-1 cells. American journal of physiology Gastrointestinal and liver physiology 282: G16–22

101. Williams GS, Boyman L, Chikando AC, Khairallah RJ, Lederer WJ (2013) Mitochondrial calcium uptake. Proceedings of the National Academy of Sciences of the United States of America 110: 10479–10486

102. Zaichick SV, McGrath KM, Caraveo G (2017) The role of Ca(2+) signaling in Parkinson’s disease. Disease models & mechanisms 10: 519–535

103. Zhao F, Li P, Chen SR, Louis CF, Fruen BR (2001) Dantrolene inhibition of ryanodine receptor Ca2+ release channels. Molecular mechanism and isoform selectivity. The Journal of biological chemistry 276: 13810–13816

104. Zhao S, Liu W, Wang J, Shi J, Sun Y, Wang W, Ning G, Liu R, Hong J (2017) Akkermansia muciniphila improves metabolic profiles by reducing inflammation in chow diet-fed mice. Journal of molecular endocrinology 58: 1–14

